# Reactivation of threat conditioning memory in humans: disentangling the effects on emotional memory and cognitive biases

**DOI:** 10.1101/2025.06.23.661086

**Authors:** Soledad Picco, Luciano Cavallino, Paula Hoijer, Juan Cruz Berón, Rodrigo S Fernández, Maria E Pedreira

## Abstract

Learning to detect and respond to threats is fundamental for survival and is often modeled through threat conditioning (TC) paradigms. While these paradigms reliably produce implicit memories that elicit physiological and behavioral responses to conditioned stimuli (CS), less is explored about how TC influences cognitive and emotional biases, particularly those implicated in anxiety disorders, such as threat overestimation and negative stimulus representation. In this study, we investigated the dynamic interaction between the reactivation of the implicit threat memory and these cognitive biases using a validated TC paradigm in humans.

In Experiment 1, participants underwent TC on Day 1, followed by a memory reactivation session (incomplete reminder: one unreinforced CS+) and a highly demanding working memory (HWM) task, used as an amnesic manipulation, or a control condition on Day 2. On Day 3, memory retention was tested using a simplified, single-trial protocol (one CS+, one CS−, and one neutral CS), followed by tasks assessing threat valuation and representation. Results indicated that the HWM task administered post-reactivation significantly reduced skin conductance responses (SCRs) and attenuated cognitive biases, without altering expectancy of the unconditioned stimulus (US).

In Experiment 2, we evaluated the effect of varying reactivation frequency (none, one, or two reminders) on implicit memory and cognitive biases. While repeated reactivations generalized the conditioned response to other stimuli, cognitive and emotional biases remained stable, suggesting a dissociation between memory generalization and evaluative processing.

These findings demonstrate that implicit threat memories can be selectively modified through post-reactivation interventions, affecting both physiological and cognitive-emotional domains. Importantly, the distinct effects of memory reactivation and reconsolidation on physiological versus cognitive outcomes support the existence of temporally and functionally dissociable mechanisms. This research highlights the need to consider cognitive biases alongside physiological responses when evaluating memory-based interventions and offers novel insight into mechanisms underlying anxiety maintenance and treatment.

## Introduction

Learning about threats in their environment is a survival behavior that allows organisms to associate previously neutral stimuli with aversive stimuli or events, generating implicit memories that trigger defensive behaviors (LeDoux & Daw, 2018; Nesse, 1998). The behaviors that are acquired serve to prevent direct harm and enhance survival. At times, these learned responses generalize to a variety of related stimuli or situations reminiscent of the aversive experience, a phenomenon known as generalization (Hull, 1943). Furthermore, individuals who struggle to inhibit their responses to threats in safe environments may find that their survival behaviors become maladaptive. This dysfunction can lead to the emergence of psychological conditions, including anxiety disorders and post-traumatic stress disorder (Dymond et al., 2015). Notably, exacerbated generalization, also called overgeneralization (Lissek & Grillon, 2015), is considered a critical pathogenic mechanism in the development of anxiety disorders. The behavioral mechanisms associated with this learning, from adaptive to dysfunctional, have been investigated using the laboratory’s threat conditioning (Pavlovian conditioning; (Pavlov, 1927) paradigms. In brief, a neutral stimulus (the conditioned stimulus or CS) is paired with an aversive unconditioned stimulus (US). Then, the CS alone elicits a conditioned response (CR) after one or a few CS-US presentations. In the case of generalization, CR is evoked by different stimuli that echo the original CS or its perceptual dimensions (i.e., perceptual generalization; (Honig & Urcuioli, 1981). Due to its significance and widespread application in the field, this paradigm is one of anxiety’s most established translational models (Boddez et al., 2014). Threat conditioning has been described from invertebrates to humans, providing essential knowledge about their physiological bases, neuronal substrates, and associated behaviors (Aguado, 2003; Krause & Domjam, 2017). In general, threat conditioning paradigms in humans analyze acquired responses through physiological changes (measured by electrodermal conductance or startle response; Kindt et al., 2009; Schiller et al., 2010). The declarative component of recognizing the stimulus associated with the aversive stimulus compared to others is also evaluated (expectancy; Kindt et al., 2009; Lovibond & Westbrook, 2024; Schiller et al., 2010).

Remarkably, this approach overlooks the intricate cognitive-behavioral effects stemming from implicit memory, neglecting its potential to influence cognitive biases associated with threat stimuli (Korteling & Toet, 2020). In this context, cognitive biases can be defined as thought patterns that frequently deviate from logical reasoning, often resulting in systematic errors or reliance on cognitive shortcuts (Fernandez et al., 2018, Leung et al., 2022, Tversky & Kahneman, 1974). Then, the affected subjective measures reflect changes in the evaluation of stimuli concerning assessing their positive or negative valence and representation understanding as the level of aversion (Grupe & Nitschke, 2013; Insel et al., 2010). These evaluations generated from the acquired association are scarcely described. To address this shortcoming, we designed an aversive-implicit memory paradigm (threat conditioning TC) that includes tasks that evaluate cognitive and emotional biases. It comprised two angry faces (CS+ and CS) and one neutral face (CSn), and the CS+ was presented with a tone (US). The subjects underwent a threat-conditioning procedure and then performed several tasks to explore standard features observed in anxiety (Bouton et al., 2001; Mineka & Zinbarg, 2006; Nader et al., 2013), such as the overestimation of threats and their consequences (Valuation) and exacerbated negative representation (Stimulus Representation). The results of this first study showed that TC produces cognitive and emotional bias 48 hours after conditioning (Fernández et al., 2018).

The flexibility of the stored information has attracted significant attention in recent decades. Essentially, presenting a reminder (a cue initially included during acquisition) serves to reactivate the previously acquired representation, necessitating a subsequent round of stabilization, known as reconsolidation. This phenomenon offers a pathway for updating consolidated memories (Dudai, 2012; Hardt et al., 2010).Consequently, a commonly suggested approach involves post-reminder interventions to modify dysfunctional memories. TCs are mainly used in non-humans and human animals to study the possibility of weakening these aversive memories. CS+ without reinforcement, incomplete reminder (Fernández et al., 2016; Pedreira et al., 2004), is generally a standard reminder in different experimental designs. Other reports have shown that this reminder can improve performance during the test session, disclosing an increase in memory precision or persistence (de Oliveira Alvares et al., 2012; De Oliveira Alvares et al., 2013; Lee, 2009).

Few studies have demonstrated the generalization associated with destabilizing the original memory. In one of them, Gazarini and colleagues (Gazarini et al., 2013) analyzed how consolidation and reconsolidation, both processes of memory stabilization, might imply a dissimilar role of the noradrenergic system. To achieve this goal, they compared the effect of yohimbine-induced noradrenergic activity during the consolidation and reconsolidation of contextual threat memory in rats. They showed that this a2-adrenoceptor antagonist potentiates memory expression by increasing the freezing response when the drug is administered immediately after acquisition or retrieval. Interestingly, the animals showed a conditioned response in both treatments when the evaluation occurred in an unpaired context, showing a generalization effect.

In a recent study using the paradigm described above (Fernandez et al., 2018), we studied the impact of attenuating the expression of TC memory using a highly demanding working memory task (HWM) after memory reactivation on physiological response (skin conductance response, SCR), expectancy, and measures of cognitive bias towards threat. The retention of conditioned threat memory was assessed on day 3 through an extinction session (twelve presentations of each CS) followed by a reinstatement test (three presentations of the US alone) and tasks targeting stimulus representation and valuation. Our findings demonstrate that the reminder-dependent intervention with HWM attenuated memory retention, as evidenced by diminished SCR and reduced representation and valuation towards the threat while leaving the expectancy of the unconditioned stimulus (US) unaffected (Picco et al., 2022).

Despite conducting assessments of cognitive and emotional biases following extinction and reinstatement sessions, we could clearly illustrate the distinct impact of implicit memory on the representation and valuation of stimuli determined by differences in implicit memory strength. However, during the testing session, one must not disregard the effect of repeated presentations of each conditioned stimulus (CS), the extinction phase, and the unconditioned stimuli (US), the reinstatement phase. These experimental manipulations could influence cognitive bias, leading to alterations or masking in their expression (Van Damme et al., 2006). Thus, to gain new insights into the dynamic relationship between implicit aversive representation and valuation cognitive processes implicated in the maintenance of anxiety features, our research took an additional step forward. In this study, we investigated the impact of TC memory on emotional and cognitive biases under various conditions. To achieve this, we pursued the following two primary objectives. First, we aimed to analyze the impact of TC memory on stimuli representation and valuation, both with and without amnesic treatment, that is, the validated HWM task. For this purpose, we conducted a test consisting of only one trial for each CS, specifically designed to exclude the influence of the extinction session. Afterward, our focus shifted to investigating the effects of presenting the incomplete reminder, designed to reactivate threat conditioning (TC) memory with no other manipulation. We aimed to go beyond assessing the direct impact on implicit memory alone and explore the potential implications of these changes associated with its reactivation and restabilization on emotional and cognitive biases influenced by implicit memory. Our working hypothesis posited that manipulating threat conditioning (TC) memory could alter cognitive and emotional biases based on dynamic interactions. We anticipated observing a fine-tuning between these processes because of the manipulations performed on implicit memory.

Therefore, in Experiment 1, we included two groups to analyze the effect of a post-reactivation treatment on TC and cognitive biases toward a threat, employing a highly demanding working memory task (HWM) as an interfering intervention and utilizing a brief testing session analyzing its effectiveness with this new simplified testing-protocol. Participants were trained on TC on Day 1. On Day 2, one group was exposed to one CS+ alone (incomplete reminder, R) and performed an HWM task, while the other group only performed the HWM task. We tested the threat-conditioning memory on Day 3 with an unreinforced presentation of each CS (short testing session: one CS+, one CS-, and one CSn). Immediately after, the subjects completed tasks targeting cognitive biases, such as stimuli representation (aversiveness) and valuation (probability and cost).

After showing that the incomplete reminder could reactivate TC memory and the amnesic effect could be demonstrated with this short testing session, the goal of Experiment 2 was to study the impact of different numbers of reactivations on implicit memory and cognitive and emotional biases. The subjects were trained on the TC task on Day 1 and divided into three groups on Day 2: one group was exposed to one R, the second group was exposed to two R, and the last group received no treatment and did not even assist to the laboratory (no day 2 group, nD2). On Day 3, we evaluated implicit memory retention, stimuli representation, and valuation using the same testing session as in Experiment 1.

Our findings suggest two primary outcomes. First, the attenuation of implicit memory expression and cognitive biases, evaluated in a simplified test session (featuring one CS+, one CS-, and one Cn), may be attributed to the impact of the HWM intervention on the shared resources of the salience and central executive networks during memory reactivation. Second, our analysis of the dynamic relationship between implicit memory reactivation and cognitive biases indicates that repeated reactivations generalize implicit memory while cognitive biases remain unaffected. We discuss this finding in terms of distinct time windows for independent processes.

## Experimental procedures

### Experiment 1

#### 1. Materials

##### 1.1 Participants

Before the experiment, all participants provided written informed consent, approved by the Ethics Committee of the Review Board of the Sociedad Argentina de Investigación Clínica (SAIC, Protocol #02/16), following the Declaration of Helsinki. A total of 44 undergraduate and graduate students from the University of Buenos Aires were included. This group consisted of 25 males and 19 females with a mean age of 23.0 (± 0.4) years. Four other subjects were excluded from the analysis; two subjects were omitted based on the exclusion criterion of the Subjective Assessment (1.2.1) and two participants were excluded due to technical issues failing to record electrodermal activity during the training session. Participants were randomly assigned to one of two groups, each comprising 22 individuals: Reactivación-HWM (R-HWM, consisting of 10 females and 12 males) and noReactivación-HWM (HWM, comprising 9 females and 13 males).

##### 1.2 Stimuli

###### 1.2.1 Subjective Assessment

The State-Trait Anxiety Inventory (STAI-S and STAI-T) (Spielberger, 1970) and the Beck Anxiety Inventory (BAI) (Beck et al., 1990) questionnaires were implemented to check the participants’ anxiety and depressive levels, as they may affect fear conditioning, especially threat conditioning (Boddez et al., 2012; Browning et al., 2015). Participants with scores higher than STAI > 45 or BAI > 30 were excluded from the analysis. Considering this criterion, two participants were excluded from the analysis as mentioned in the previous section (1.1)

###### 1.2.2 Threat Conditioning

Conditioned stimulus. Three distinct male facial images from the Karolinska Directed Emotional Faces database were used as conditioned stimuli (CS). These images were displayed centrally on a black screen (each slide measuring 9.5 cm × 7 cm). Two represented anger expressions (CS+ and CS-), while the third remained neutral (CSn). The aversive images used as CS+ and CS-, were counterbalanced across participants. See FIgure 1 for the faces implemented.

**Figure 1.**
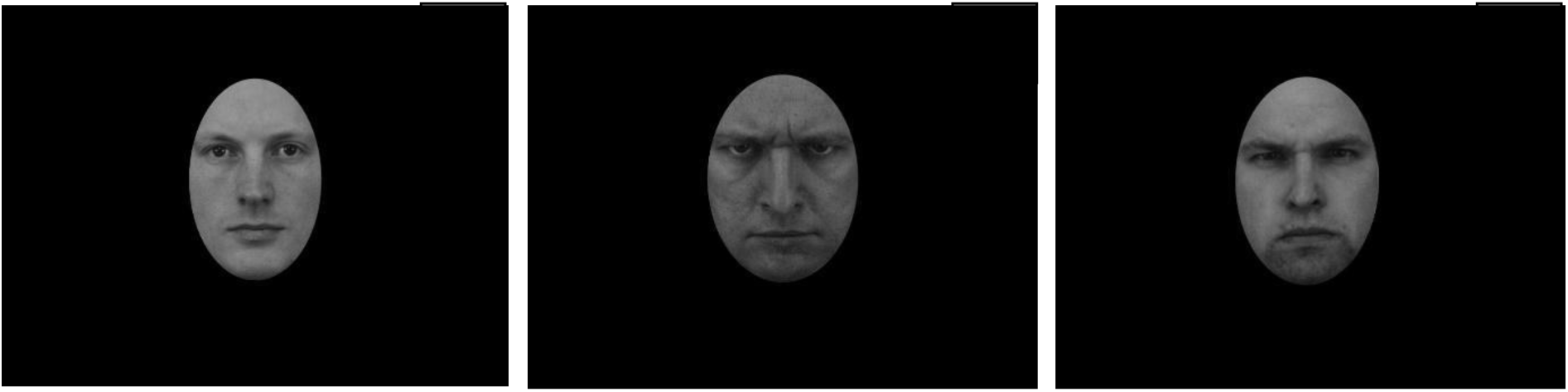
(1) Neutral face, CSn; (2) The anger-expressed face with tone, CS+; and (3) anger-expressed face without tone CS−.

Unconditioned stimulus. The unconditioned stimulus (US) consisted of an auditory tone lasting 1.5 seconds, delivered through stereo headphones. This tone was generated by a TG/WN Tone-Noise Generator (Psychlab) and digitally controlled to maintain a mean of 98 dB ± 2 dB. The intensity of the US was individually adjusted for each participant to evoke sensations described as “unpleasant but not painful.” This adjustment involved selecting a decibel level that elicited self-reported aversion to the US. The same intensity level of the US was consistently applied to each participant throughout the reinstatement phase.

###### 1.2.3 Working Memory Intervention

The Pace Auditory Serial Addition Test (PASAT; (Gronwall, 1977) is a complex cognitive task that encompasses various cognitive functions, particularly those associated with attention, and is recognized as a measure of working memory (WM) proficiency (Range, 2023). Hence, in this study, PASAT was employed as the high-demanding working memory task (HWM). Participants were presented with a prerecorded series of numbers ranging from 1 to 9 via headphones and instructed to add the last two numbers of each sequence. The task consisted of 120 trials, with numbers presented at 2-second intervals (PASAT 200 block a + b, (Gronwall, 1977). Participants were not provided with feedback; only responses within the stipulated interval were deemed correct. For instance, if the sequence presented were ‘2’, ‘4’, and ‘8’, the correct responses would be ‘6’ followed by ‘12’.

To assess working memory capacity, the initial 60 trials of the PASAT (Gronwall, 1977) were employed. These were scrutinized using arousal and performance metrics. Specifically, arousal levels were evaluated using Skin Conductance Level (SCL), quantified as the mean tonic signal (MS) recorded throughout the task. Furthermore, the accuracy rate of responses served as an indicator of working memory performance.

###### 1.2.4 Cognitive biases assessment

Stimulus representation. Subjects rated the aversiveness of 12 different pictures of male faces using a 0–8 scale (five angry, four neutrals, and the ones used as CS+, CS-, and CSn). Pictures were randomly presented on the screen until a response was given with the keyboard. The generalization score is for all aversive faces (excluding the CS+ and CS-) and for all neutral faces (excluding the CSn).

Valuation (Probability and Cost). Participants were presented with a total of 96 images corresponding to the CS+ and CS-stimuli, with the objective of eliciting assessments of positive and negative probability and cost scores. 24 of these images had positive scenarios and 24 negative scenarios, each involving the corresponding CS+ or the CS-image. In response to each scenario and associated CS, participants initially rated the perceived likelihood of the hypothetical event on a 0–8 scale. Subsequently, participants rated the perceived desirability or averseness of the scenario using the same 0–8 scale. For instance, a negative scenario prompted questions such as, “How likely would it be for HIM to talk bad about you at work?” followed by “How bad would it be for HIM to talk bad about you at work?”. Conversely, a positive scenario included inquiries like, “What is the likelihood that you tell a joke, and HE laughs?” followed by “How much would you like it if you tell a joke and HE laughs?”.

#### 2. Measurements

Implicit memory. SCR was employed as the dependent variable and key physiological indicator throughout the experiment to gauge the acquisition and retention of threat-conditioning (Knight et al., 2010; Oyarzún et al., 2012; Schiller et al., 2010; Zimmermann & Bach, 2020). Through the presentation of CS faces and US tones, the conditioned response (CR) evaluation involves measuring SCR to discern underlying mechanisms of unconscious aversive learning (Stemerding et al., 2022). Electrodermal activity was recorded using an input device (Psychlab Precision Contact Instruments) featuring a sine excitation voltage (±0.5 V) at 50 Hz. Two Ag/AgCl electrodes (20 mm × 16 mm) were positioned on the intermediate phalanges of the index and middle fingers of the non-dominant hand.

Explicit memory. The evaluation of declarative memory in the context of TC involved the utilization of an external keyboard with Yes/No buttons. Subjects were tasked with indicating their expectancy of the US during each CS presentation. Specifically, participants were instructed to press the ‘YES’ button if they anticipated the occurrence of the US (tone) following the CS or the ‘NO’ button if they did not expect the US. This task was performed across all phases of threat conditioning, encompassing acquisition, reactivation, and reinstatement.

#### 3. Procedure

All participants completed the STAI and BAI questionnaires (1.2.1) online during the experimental week. Initially, the participants were randomly divided into experimental groups; Reactivation-HWM or noReactivation-HWM. On the first day, participants had to sign the informed consent form and read the experimental procedure. There was an experimental training session to ensure the participants understood the experiment. Habituation trials showed the CS+, CS-, and CSn three times in a random order to familiarize the participants with the faces used in the experiment. The experimental procedure started with the Stimuli Representation followed by Acquisition phase (3.1), where the participants underwent threat conditioning. Following, the Cognitive Bias Assessment Stimulus representation (1.2.3) was administered. On the second day, within the Reactivation phase (3.2), the Reminder-HWM group received a stimuli reminder (CS+). Both groups had to perform the HWM task. The memory Evaluation phase (2.3) on the last day was performed to test the retention of threat-conditioning memory. This test was followed by the Cognitive Biases Assessments: Stimulus Representation and Valuation (Probability and Costs) (1.2.4). The experimental days were consecutive; see Figure 2 for an overview.

**Figure 2.**
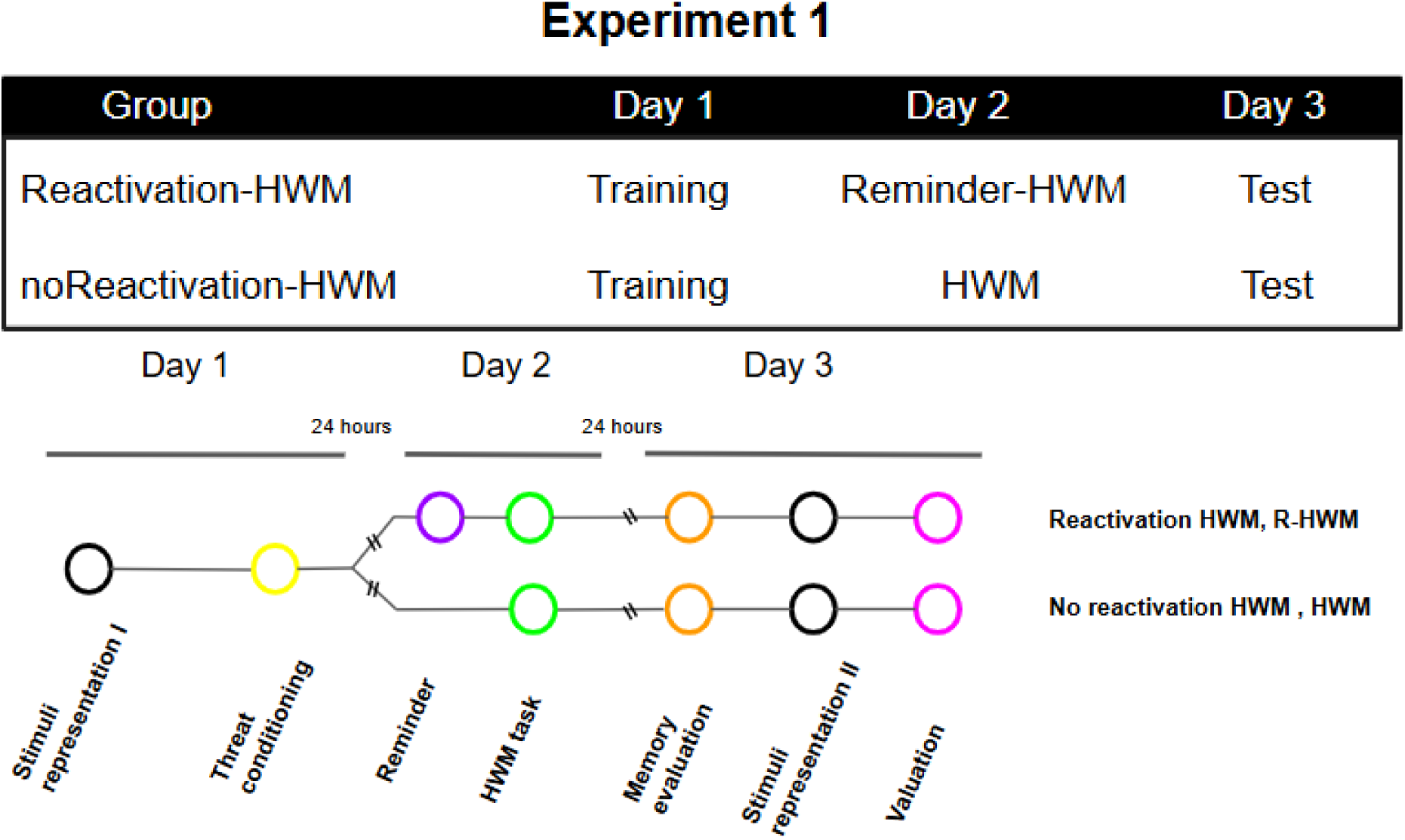
Schematic depiction of the experimental design illustrating group specifications throughout the three experimental sessions. Chronological outline delineating the experimental schedule for the two groups: those undergoing either no-reactivation or reactivation concurrently with a Hard Working Memory (HWM) task.

##### 3.1 Acquisition Phase (Day 1)

On the first experimental day, the participants underwent a TC. Each face stimulus was presented eight times, with 24 trials in total for the three different faces. The face-CS+ was associated with a tone-US in 75% of its presentations since violations of threat expectancy are essential for threat conditioning (Willems & Vervliet, 2021). The unconditioned face images (CS− and CSn) were not paired with the tone-US. The CS was presented for 6 seconds, and the US onset occurred 4.5 seconds after CS onset, overlapping with the final 1.5 seconds of the CS. The inter-trial interval was variable, ranging between 8 and 12 seconds.

##### 3.2 Reactivation Phase (Day 2)

On the second day, the Reactivation-HWM group was presented with the unreinforced CS+ to reactivate their memory. Immediately after, an interruption and a black screen indicate the end of the experiment. Subsequently, both groups (R-HWM and HWM) performed the PASAT HWM task (1.2.3). The interval separating the reminder and HWM tasks was 5 minutes after the CS+ presentation.

##### 3.3 Evaluation Phase (Day 3)

On the final day of the experiment, all participants were exposed once to the three distinct conditioned stimuli types (CS+, CS-, and CSn). The presentation duration for each stimulus was consistent at 6 seconds, with inter-trial intervals ranging from 8 to 12 seconds. Notably, no US was presented during this phase. SCR and the expectancy of the US were measured as outcome variables.

### Experiment 2

#### 4. Materials

##### 4.1 Participants

As in experiment 1, all participants provided written informed consent, approved by the Ethics Committee of the Review Board of the Sociedad Argentina de Investigación Clínica (SAIC, Protocol #02/16) following the Declaration of Helsinki. A total, of 70 undergraduate and graduate students from the University of Buenos Aires participated (34 females, 36 males; mean age 23.2 ± 0.4 years). Twelve participants were excluded: two due to their scores and the subjective assessment (1.2.1) and ten due to equipment failures. Participants were assigned to three groups: ND2 (control, no treatment on Day 2, n = 21), 1-R (one CS+ reminder, n = 23), and 2-R (two CS+ reminders, n = 26).

##### 4.2 Stimuli

Stimuli were identical to those used in Experiment 1, see 1.2.1 Subjective assessment, 1.2.2 Threat conditioning and 1.2.4 Cognitive bias assessment. Participants with scores higher than STAI > 45 or BAI > 30 were excluded from the analysis. Considering this criterion, two participants were excluded from the analysis as mentioned in the previous section (1.1)

#### 5. Measurements

The measurements, of both, implicit and explicit memory are conducted similarly to Experiment 1.

#### 6. Procedure

The phases of the experiment are explained in Experiment 1. Differences in the reactivation phase on day 2 were explained in 6.2 Reactivation Phase. See Figure 3 for an overview

**Figure 3.**
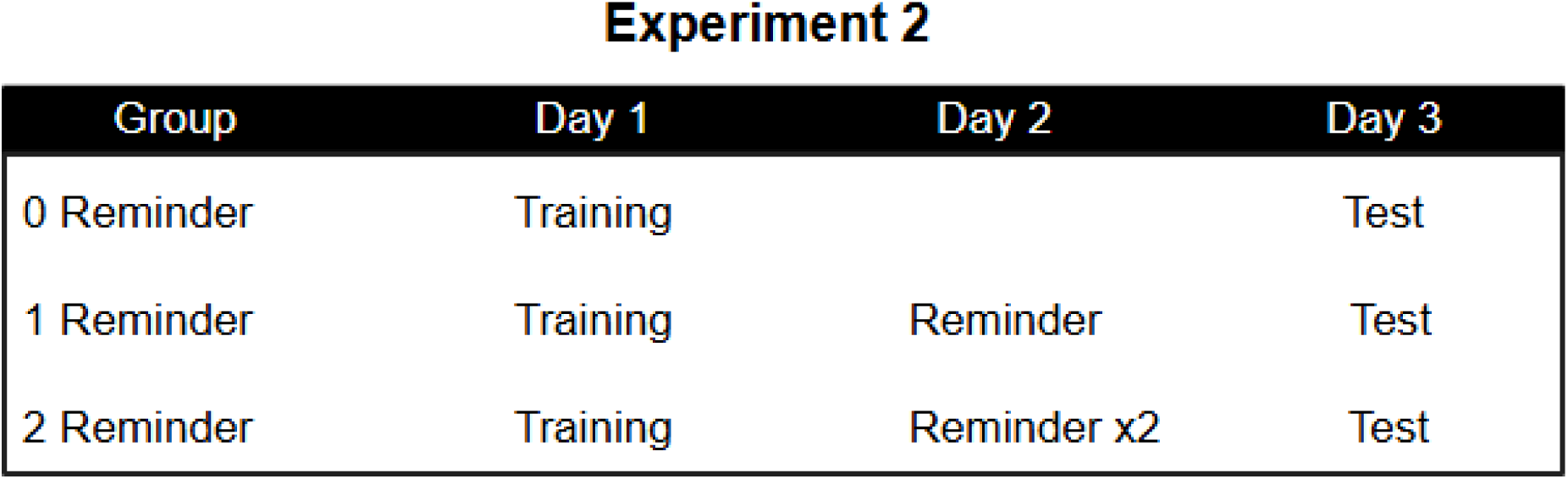
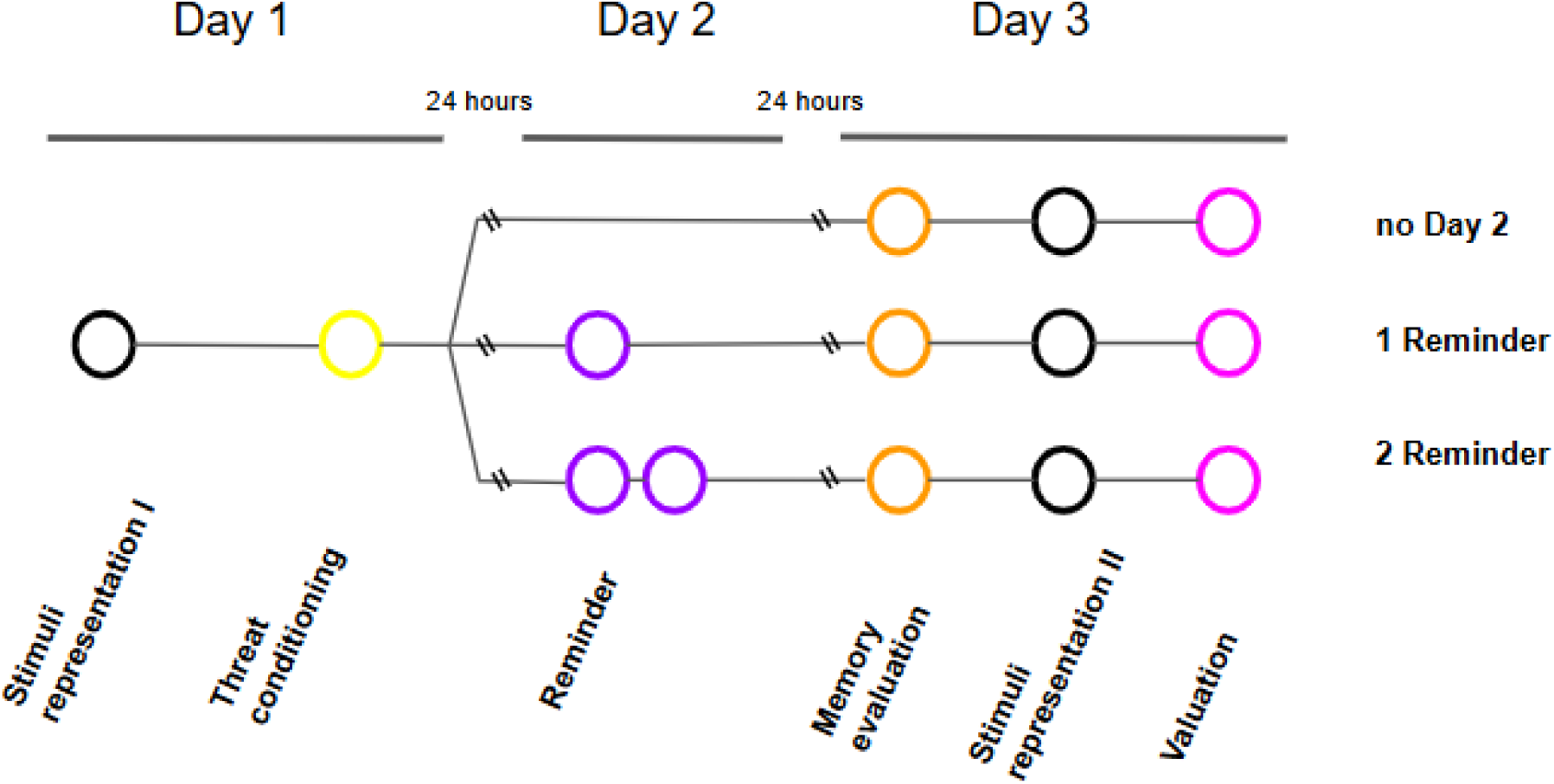
Schematic depiction of the experimental design illustrating group specifications throughout the three experimental sessions. Chronological representation illustrating the experimental schedule for the three groups categorized by the absence of a reminder, presence of a singular reminder, or exposure to two reminders.

##### 6.1 Acquisition Phase (Day 1)

Threat conditioning was conducted identically to Experiment 1.

##### 6.2 Reactivation Phase (Day 2)

During the reactivation phase, two groups (1-R and 2-R) received a presentation of the unreinforced CS+. Where 1-R was presented with one stimulus reminder (CS+), the 2-R group saw two reminders of the CS+ faces consecutively with a short interruption of 2 seconds in between. No US (tone) was presented during the presentation. Instructions were similar on Day 1, so people expected to see the entire experiment. The control group, nD2, does not participate in the experiment on day 2.

##### 6.3 Evaluation Phase (Day 3)

The evaluation phase was conducted in the same way as Experiment 1.

#### 7. Statistical Analysis

Subjective assessments were analyzed using a t-test (Experiment 1) and a One-Way Analysis of Variance, ANOVA (Experiment 2). Due to the considerable variation in individual sensitivity to SCR across different tasks and participants, the resultant SCR values were standardized across conditions. Therefore, it involved converting each time point into z-scores, achieved by subtracting the mean and dividing by the standard deviation of the two conditions, as detailed in previous studies (Ben-Shakhar, 1985; Mas-Herrero et al., 2014). Subsequently, a five-point average moving filter (applied through the “smooth” command in the Matlab software) was employed to enhance the smoothness of the SCR raw signal. The designated time window for signal extraction ranged from 0.5 to 4.5 seconds. SCR measurements during threat-conditioning were analyzed using a robust Linear Mixed Model, also known as a hierarchical or multilevel model. Separate linear mixed models were employed to examine the effects of fixed factors (Group, Stimulus, Trial) on various memory processes, including memory acquisition, retention, and evaluation. Planned comparisons were conducted for specific trial pairs within each memory process. All analyses incorporated Subject-level intercepts as random effects (SCR ∼ Group * CS * Time + (1| ID)). The significance of fixed factors was assessed using the Likelihood-Ratio test, with Satterthwaite’s approximation utilized for degrees of freedom. Using R Studio, mixed models were implemented (using the lme4 package), and planned comparisons were conducted using the Tukey method (using the emmeans package) with the Kenward-Roger approximation for degrees of freedom. Similarly, cognitive biases (Stimuli representation, Valuation, and Attentional biases) were examined through Linear Mixed Models and subsequent planned comparisons, following the electrodermal activity analysis framework (i.e., Score ∼ Condition * Group + (1| ID)). Effect sizes were reported as Marginal and Conditional R2 values, with Marginal R2 indicating variance explained by fixed factors and Conditional R2 representing the proportion of variance explained by the entire model (including both fixed and random factors). R2 statistical measures were calculated using the sjstatspackage in R. The odds of responding “YES” to US expectancy during threat-conditioning and memory evaluation were analyzed using a Logistic Generalized Linear Mixed Model with Trial as a repeated factor. Finally, electrodermal activity (SCR) and the proportion of correct responses were analyzed using ANOVA, followed by planned comparisons with Tukey adjustment for multiple comparisons.

## Results

### Experiment 1: Effect of a HWM after memory reactivation

To analyze the effect of a high-demand working memory task (HWM) on cognitive biases and memory, we evaluated the implicit and declarative components of memory, as well as the stimulus representation and valuation in the two experimental groups mentioned in section 2 (Reactivation-HWM: R-HWM, and noReactivation-HWM: HWM).

#### Subjective assessment

When analyzing the results obtained for the State-Trait Anxiety Inventory (STAI-T, STAI-S) and the Beck Anxiety Inventory (BAI), used to rule out an anxious psychopathology, it was observed that the participants did not show differences in self-reported anxiety (STAI-T, mean of the R-HWM group: 30.26 ± 2.84, HWM: 28.85 ± 2.; F2=0.150, p=0.70; STAI-S, mean of the R-HWM group: 36.33 ± 2.99, HWM: 38.38 ± 2.14; F2=0.293, p=0.59; BAI, mean of the R-HWM group: 11.40 ± 1.92, HWM: 12.75 ± 2.38; F2=0.199, p=0.66).

#### Threat memory acquisition

When analyzing the first and last trials, a significant main effect of stimulus was found, X^2^(2,194) = 5.69, p = 0.003. There was no main effect of trial, X^2^(1,195) = 0.027, p = 0.868, or group, X^2^(1,40) = 0.013, p = 0.9. Nevertheless, a significant interaction between stimuli (CSs) and trial emerged, X^2^(2,194) = 8.63, p = 0.0002, while no interaction was observed between group and trial, X^2^(1,195) = 0.028, p = 0.86. Planned comparisons showed successful conditioning for both groups, as evidenced by a greater response to CS+ than to CS− and CSn in both groups during the last trial. Planned comparison CS+ vs. CS−: t(197) = 3.66, Estimate = 0.656, SE = 0.179, p = 0.0009. Planned comparison CS+ vs. CSn: t(197) = 4.39, Estimate = 0.794, SE = 0.181, p = 0.0001. No differences were found between CS− and CSn: t(196) = 0.756, Estimate = 0.137, SE = 0.182, p = 0.73. Figure 4. All remaining interactions are presented in the supplementary table 1a.

**Figure 4.**
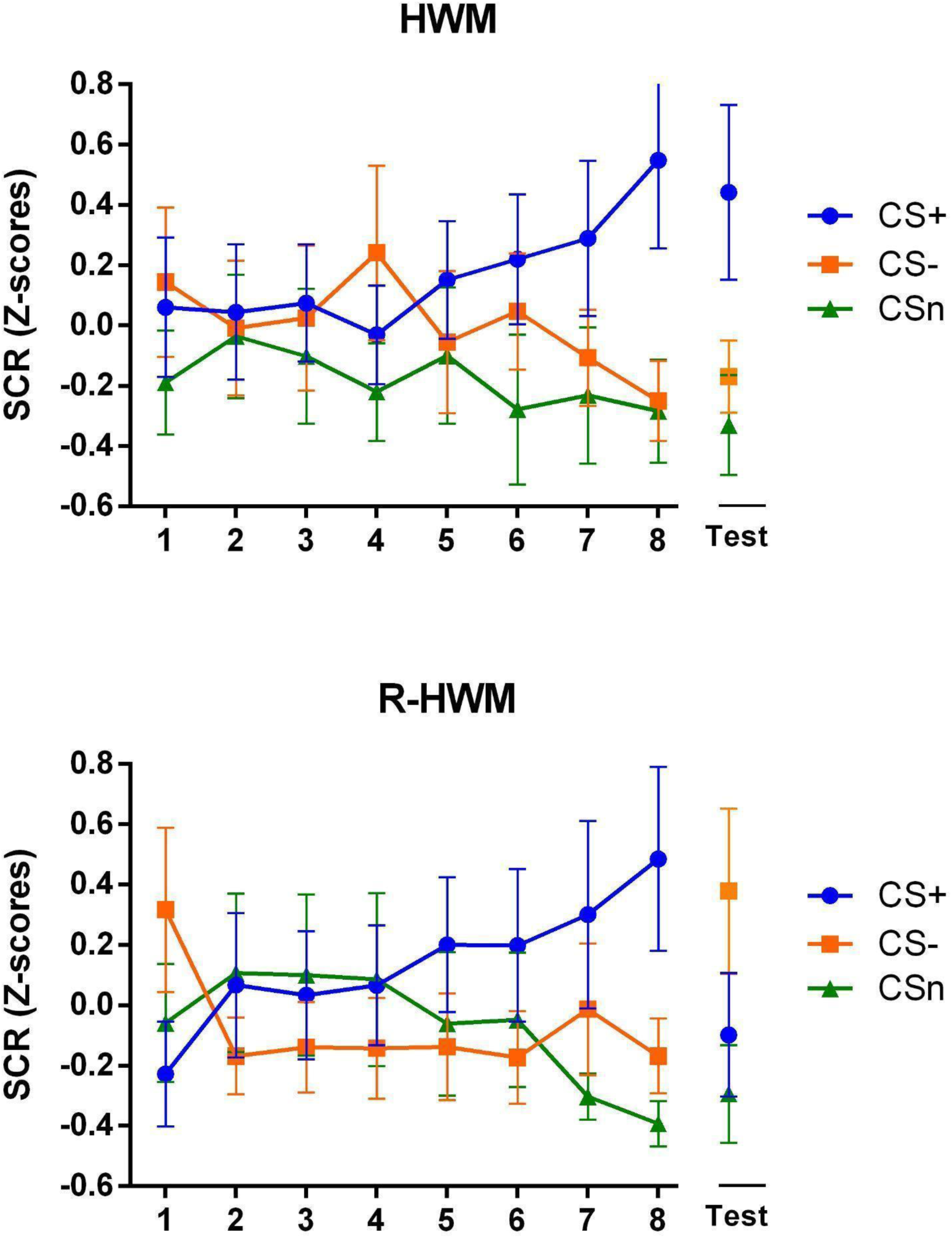
Reconsolidation interference of Threat Memory. (A) Reactivation-HWM group (R-HWM). (B) noReactivation-HWM group (HWM). Mean SCR (Z-score) ± SEM for the eight trials of Day 1 (Threat conditioning acquisition) and the Test of Day 3.

#### Threat memory retention

When comparing the last trial of Day 1 with the test trial (day 3), a significant main effect of stimulus was found X^2^(2,190)=11.33, p=0.00002. There was no main effect of trial, X^2^(1,196) = 0.018, p = 0.89, or group, X^2^(1,41) = 0.0001, p = 0.9. Nevertheless, a clear trend was emerged indicating that stimuli changed differently depending on the group and time. Group × Stimulus interaction: X^2^(2,190)=2.61, p=0.07. Time × Stimulus interaction: X^2^(2,190)=2.57, p=0.07. Planned comparisons were conducted to assess how stimuli changed for each group. While the CS+ did not change between the last trial of Day 1 and the test trial for the HWM group, t(190)=0.324, Estimate=0.087, SE=0.27, p=0.746, it did decrease in the R-HWM group during the test trial, t(191)=2.17, Estimate=0.58, SE=0.27, p=0.03. This pattern highlights the impact of a demanding task on memory retention after a reminder presentation. Figure 4. All remaining interactions and comparison are presented in the supplementary table 1b,c,d.

Furthermore, when comparing the three stimuli on Day 3 for each group, differences were observed between CS+ and both CS− and CSn in the HWM group: planned comparisons CS+ vs. CS-: t(188)=2.33, Estimate=0.611, SE=0.262, p=0.054; CS+ vs. CSn: t(192)=2.706, Estimate=0.752, SE=0.278, p=0.02; CS− vs. CSn: t(192)=0.507, Estimate=0.1410, SE=0.278, p=0.868. However, in the reactivated group, R-HWM, no differences were found between CS+ and either CS− or CSn: planned comparisons CS+ vs. CS-: t(188)=-1.73, Estimate=-0.476, SE=0.275, p=0.196; CS+ vs. CSn: t(189)=0.731, Estimate=0.204, SE=0.279, p=0.745; CS− vs. CSn: t(189)=2.438, Estimate=0.68, SE=0.279, p=0.041. This result indicates interference with memory retention when an HWM is performed after a reminder on Day 2. Figure 4

#### Unconditioned stimulus expectancy

While differences in implicit memory were observed between groups, no distinctions were found in the declarative component of this memory. These findings are consistent with previous results (Picco et al., 2022). During acquisition, both groups exhibited higher odds of responding ‘YES’ after the CS+ compared to the CS-(Odds Ratio CS+ vs. CS-= 0.54, CI 95% [0.36-0.71], p < 0.001) or the CSn (Odds Ratio CS+ vs. CSn = 0.82, CI 95% [0.61-1.4], p < 0.001). The US expectancy associated with CS+ remained consistent from the end of threat acquisition (Day 1) to the beginning of the retention trial (Day 3) (Odds Ratio CS+ vs. CS-= 0.67, CI 95% [0.57-0.78], p < 0.001; Odds Ratio CS+ vs. CSn = 0.88, CI 95% [0.78-0.99], p < 0.001). Furthermore, these differences persisted in the retention trial on Day 3 (Odds Ratio CS+ vs CS-= 0.66, CI 95% [0.51-0.81], p < 0.001; Odds Ratio CS+ vs. CSn = 0.92, CI 95% [0.77-1.06], p < 0.001). Supplementary material figure 1.

#### Working memory intervention

To evaluate the cognitive demanding nature of the HWM, in previous work, the SCR and performance were compared between the high-demanding WM task (HWM) and a low-demanding WM task (LWM) (Picco et al., 2022). The higher SCL and lower performance of the HWM confirmed the demanding nature of the task (Picco et al., 2022). In the present work, the same task (HWM) was used. Therefore, SCR and performance were compared to evaluate the similarity between the experimental groups (R-HWM and HWM). No differences were found for the SCR (Group F (1.15) = 0.22, p = 0.65, fixed-effects correlation ηp^2^ = −0.598; Planned comparison, Reactivation-HWM vs noReactivation-HWM, t(16) = 0.47, p = 0.65, Estimate=-0.22, SE= 0.35), nor for performance (Group F (1.17) = 0.02, p = 0.89, ηp^2^ = −0.628; Planned comparison, R-HWM vs HWM, t(16) = −0.13, p = 0.89, Estimate-14, SE= 13.0) showing no differences between groups.

#### Cognitive biases

##### Stimulus representation

Before conditioning (Day 1), no differences were found between groups in the levels of aversiveness for the CS+ (Planned comparison for CS+ Stimulus, Day 1 HWM vs. R-HWM: t(53) = −0.30, p = 0.76, Estimate=-0.22, SE=0.73) or CS-(Planned comparison for CS-Stimulus, Day 1 HWM vs. R-HWM: t(53) = 0.11, p = 0.91, Estimate=0.08, SE=0.73).

Differences in the stimulus representation between Day 1, before training, and Day 3, following the retention test, were observed (Main effects: Stimulus X^2^ (1,90) = 1.65, p = 0.2, Group χ^2^ (1,90) = 0.05, p = 0.82, Time X^2^ (1,90) = 7.87, p = 0.006; Conditional R^2^ = 0.660, Marginal R^2^ = 0.036). Specifically, the HWM group rated the CS+ as more aversive or unpleasant on the third day than on Day 1. In contrast, the R-HWM group showed no differences (Planned comparison for CS+, Day 1 vs. Day 3, HWM: t (90) = −2.93, p = 0.004, Estimate=-1.23, SE=0.42; R-HWM: t (90) = −1.19, p = 0.24, Estimate=-0.53, SE=0.45). For the CS-, no differences were found between Day 1 and Day 3 for either of the two groups (Planned comparison for CS-, Day 1 vs. Day 3, HWM: t (90) = −9.8, p = 0.33, Estimate=-0.41, SE=0.42; R-HWM: t (90) = −0.59, p = 0.55, Estimate=-0.26, SE=0.45) Figure 5a.

**Figure 5.**
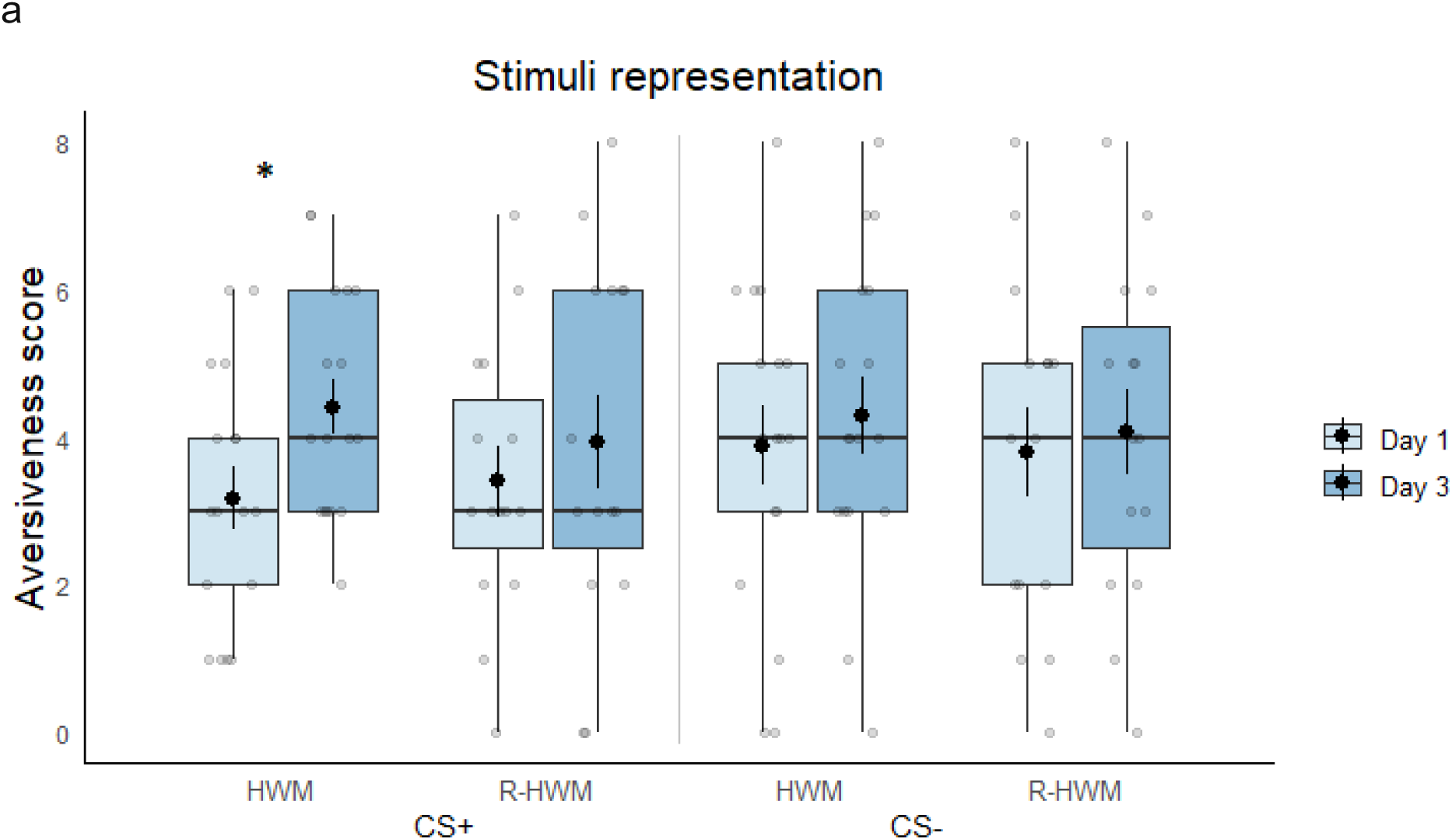
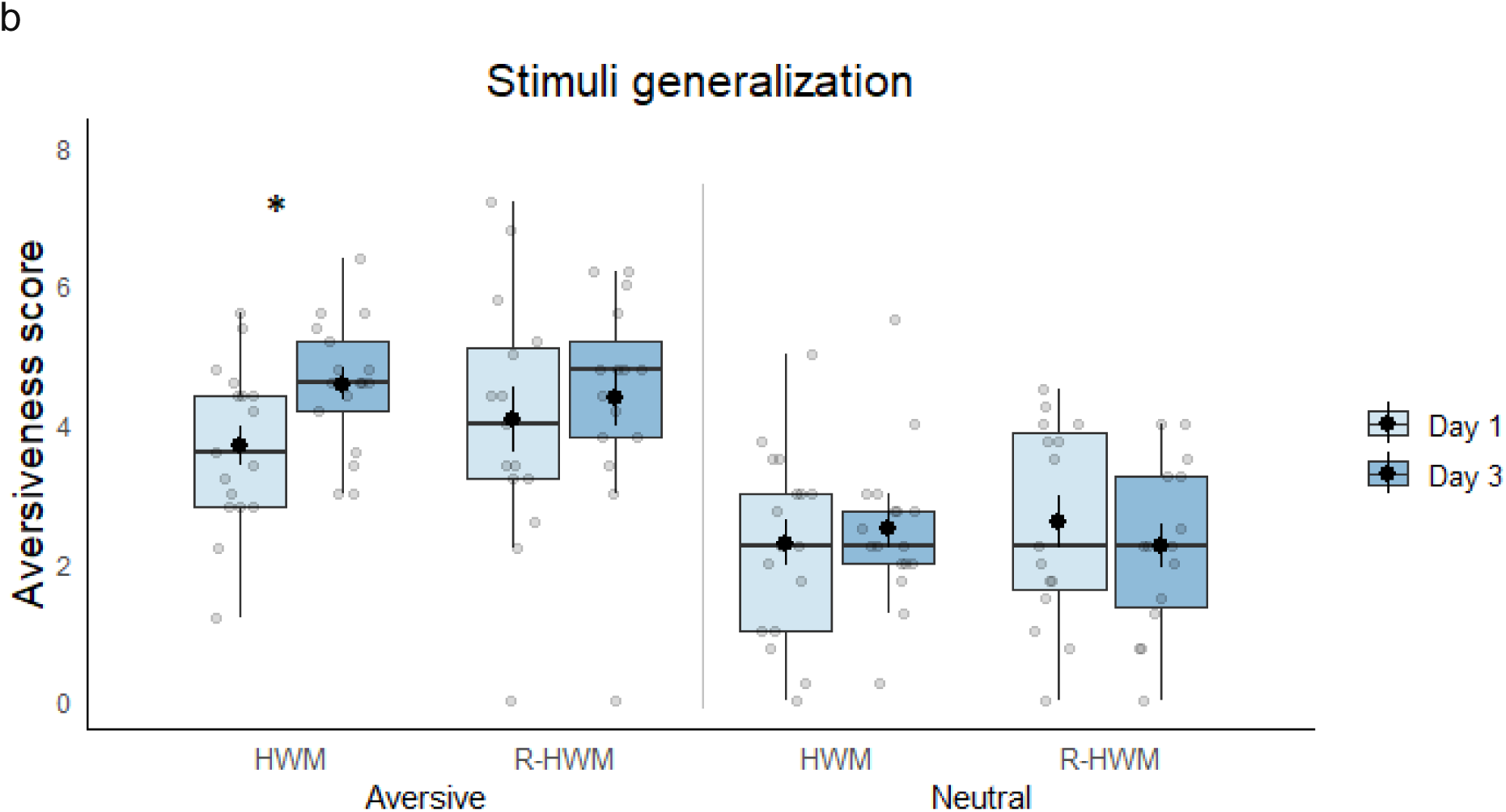
Cognitive biases assessment before threat conditioning (Day 1) and after test (Day 3). (A) Stimulus Representation (aversiveness). Aversiveness score for the CS+ and CS-in Day 1 and Day 3 for the HWM and R-HWM groups. The box plots indicate interquartile range (IQR), band inside the box represents the median, and whiskers are the maximum and minimum values. Grey circles represent the value of each individual and larger dark circle the mean. (B) Stimulus Generalization. Aversiveness score for the aversive and neutral pictures in Day 1 and Day 3 for the HWM and R-HWM groups. The box plots indicate interquartile range (IQR), band inside the box represents the median, and whiskers are the maximum and minimum values. Grey circles represent the value of each individual and larger dark circle the mean. Asterisks (*) represent significant differences between encounters, with α=0.05

Furthermore, we also separately analyzed the two groups’ aversive and neutral faces for Day 1 and Day 3, excluding the stimuli used in threat conditioning, such as CS+, CS-, and CSn. The aversive faces were scored as more unpleasant on Day 3 compared with Day 1 in the HWM group. Still, no differences were found for the R-HWM group (Main Effects: Valence X^2^ (1,538) = 147.5, p < 0.0001, Group X^2^ (1,30) = 0.02, p = 0.88, Time X^2^ (1,538) = 3.33, p = 0.07; Conditional R^2^ = 0.384, Marginal R^2^ = 0.174; Planned Comparison for Aversive Stimuli, Day 1 vs Day 3, HWM group: t (538) = −3.29, p = 0.001, Estimate=-0.88, SE=0.27; R-HWM group: t (538) = −1.17, p = 0.24, Estimate=-0.33, SE=0.29). However, no differences were found between days when neutral faces were evaluated for either of the two groups (Planned Comparison for Neutral Stimuli, Day 1 vs Day 3, HWM group: t (538) = −0.69, p = 0.49, Estimate=-0.21,SE=0.30; R-HWM group: t (538) = 1.09, p = 0.27, Estimate=0.35, SE=0.31). Figure 5b

In Day 1 and Day 3, aversive faces were scored significantly more aversive or unpleasant than the neutral faces in both groups (Planned comparison Aversive vs. Neutral Stimuli, Day 1, HWM group: t (538) = 4.98, p < 0.0001, Estimate=1.41, SE=0.28; R-HWM group: t (538) = 4.86, p < 0.0001, Estimate=1.47, SE=0.30; Day 3, HWM group: t (538) = 7.36, p < 0.0001, Estimate=2.09, SE=0.28; R-HWM group: t (538) = 7.12, p < 0.0001, Estimate=2.15, SE=0.30). Supplementary material figure 2

##### Valuation (probability and cost)

The probability of negative scenarios were higher for the CS+ than the CS− for both group, while no differences were found in the probability of positive scenarios (Main Effects: Stimulus χ^2^ (1,1392) = 4.77, p = 0.03; Group X^2^ (1,27) = 0.86, p = 0.36; Valence X^2^ (1,1392) = 61.5, p < 0.0001; Stimulus x Valence Interaction X^2^ (2,1392) = 26.2, p < 0.0001; Conditional R^2^ = 0.139, Marginal R^2^ = 0.061; Planned Comparisons, Negative Valence Probability, HWM CS+ vs. CS− t (1393) = 5.03, p < 0.0001, Estimate=1.2. SE=0.23; R-HWM CS+ vs. CS− t (1393) = 2.28, p = 0.02, Estimate=0.52, SE=0.23; Positive Valence Probability, HWM CS+ vs. CS− t (1393) = −1.53, p = 0.13, Estimate=-0.35, SE=0.23; R-HWM CS+ vs. CS− t (1393) = −1.41, p = 0.15, Estimate=-0.32, SE=0.23). Figure 6a

**Figure 6.**
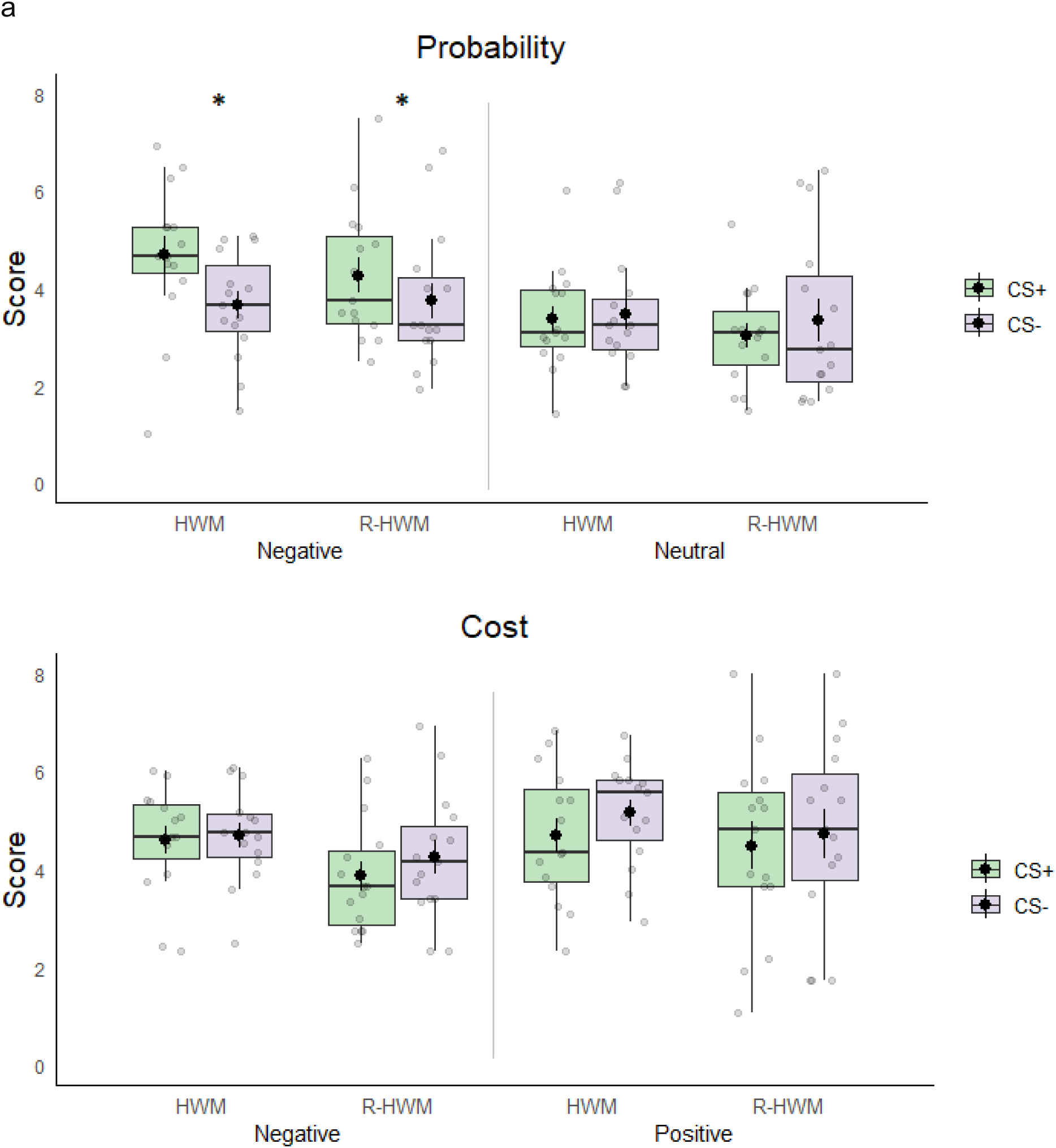
Valuation (Probability and Cost) after test (Day 3). (A) Probability: Score for Negative and positive scenarios for CS+ and CS− in the HWM and R-HWM group. The boxplots indicate interquartile range (IQR), the band inside the box represents the median, and the whiskers are the maximum and minimum values. Grey circles represent the value of each individual, and larger dark circles represent the mean. Asterisks (*) represent significant differences between encounters, with α=0.05. (B) Cost: Score for Negative and positive scenarios for CS+ and CS− in the HWM and R-HWM groups. The boxplots indicate interquartile range (IQR), the band inside the box represents the median, and the whiskers are the maximum and minimum values. Grey circles represent the value of each individual, and larger dark circles represent the mean.

When participants rated how good or bad the hypothetical situation was (positive and negative cost), no differences were found between CS+ and CS− for either of the groups (Main Effects: Stimulus χ^2^ (1,1393) = 4.78, p = 0.02; Group X^2^ (1,28) = 1.6, p = 0.21; Valence X^2^ (1,1393) = 10.17, p = 0.001; Conditional R^2^ = 0.125, Marginal R^2^ = 0.016; Stimulus x Group Interaction X^2^ (1,1393) = 0.05, p = 0.81; Planned Comparisons, Cost with Negative Valence, R-HWM CS+ vs. CS− t (1393) = −1.5, p = 0.13, Estimate=-0.39, SE=0.27; HWM CS+ vs. CS− t (1393) = −0.16, p = 0.87, Estimate=-0.04, SE=0.27; Cost with Positive Valence, R-HWM CS+ vs. CS− t (1393) = −0.94, p = 0.35, Estimate=-0.25, SE=0.27; HWM CS+ vs. CS− t (1393) = −1.78, p = 0.08, Estimate=-0.48, SE=0.27). Figure 6b

### Experiment 2: Effect of different number of incomplete reminders (R)on memory reactivation

To assess the impact of varying numbers of reminders on reactivation and memory consolidation, we evaluated implicit and declarative memory, as well as stimulus representation and valuation, in three groups: no reminder (nD2), one reminder (1-R), or two reminders (2-R).

#### Subjective assessment

When analyzing the results obtained for the State-Trait Anxiety Inventory (STAI-T, STAI-S) and the Beck Anxiety Inventory (BAI), used to rule out anxious psychopathology, it was observed that the participants did not show differences in self-reported anxiety (STAI-T, mean of group nD2: 28.08 ± 2.36, 1-R: 28.56 ± 1.82, 2-R: 30.04 ± 2.41; F_2_=0.191, p=0.83; STAI-S, mean of group nD2: 39.42 ± 1.72, 1-R: 35.69 ± 2.49, 2-R: 36.19 ± 1.71; F_2_=0.191, p=0.83; BAI, mean of group nD2: 13.25 ± 1.94, 1-R: 10.56 ± 1.64, 2-R: 13.92 ± 1.61; F_2_=1.037, p=0.36).

#### Threat memory acquisition

When analyzing the first and last trials, a significant main effect of stimulus was found, X^2^(2,307) = 7.96, p = 0.0004. There was no main effect of trial, X^2^(1,308) = 0.008, p = 0.92, or group, X^2^(2,83) = 0.039, p = 0.96. Nevertheless, a significant interaction between stimulus (CSs) and trial emerged, X^2^(2,307) = 12.9, p = 0.000004, while no interaction was observed between group and trial, X^2^(2,308) = 0.022, p = 0.977. Planned comparisons showed successful conditioning in all three groups, as evidenced by a greater response to CS+ than to CS− and CSn during the last trial. Planned comparison CS+ vs. CS−: t(310) = 4.4, Estimate = 0.632, SE = 0.144, p < 0.0001. Planned comparison CS+ vs. CSn: t(308) = 5.06, Estimate = 0.709, SE = 0.14, p < 0.0001. No differences were found between CS− and CSn: t(310) = 0.533, Estimate = 0.077, SE = 0.144, p = 0.85. Figure 7. All remaining interactions are presented in Supplementary Table 1e.

**Figure 7.**
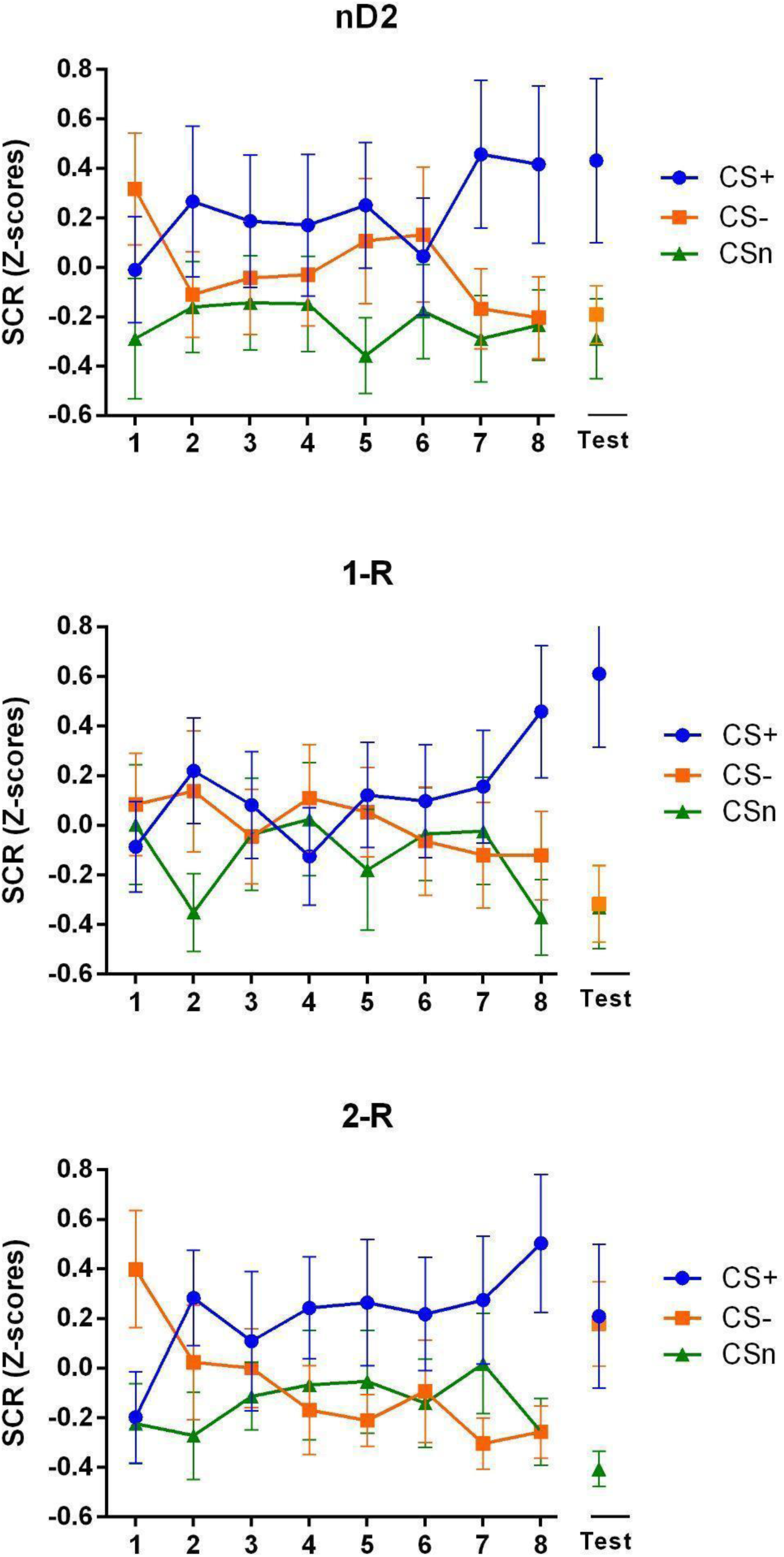
Effect of different number of reminders on threat Memory. (A) No reminder (nD2). (B) One reminder (1-R). (C) Two reminders (2-R). Mean SCR (Z-score) ± SEM for the eight trials of Day 1 (Threat conditioning acquisition) and the Test of Day 3.

#### Threat memory retention

When comparing stimuli within each group on the testing day (day 3), a significant main effect of stimulus was found X^2^(2,104)=16.1, p=0.0000007. There was no main effect of group, X^2^(2,70) = 0.018, p = 0.98. Nevertheless we found an interaction effect between group and stimulus, X^2^(4,104) = 2.43, p = 0.051.

Planned comparisons revealed that the CS+ elicited a stronger response than both CS− and CSn only in groups nD2 and 1-R. Planned comparison, Group nD2, contrast CS+ vs. CSn, t(106) = 2.55, Estimate = 0.62, SE = 0.24, p = 0.03; and a strong trend in the same direction for the contrast CS+ vs. CS-, t(108) = 2.22, Estimate = 0.56, SE = 0.25, p = 0.07; contrast CS− vs. CSn, t(110) = 0.22, Estimate = 0.05, SE = 0.26, p = 0.97. Group 1-R, contrast CS+ vs. CSn, t(106) = 3.99, Estimate = 0.99, SE = 2.5, p = 0.0004; contrast CS+ vs. CS-, t(105) = 3.84, Estimate = 0.92, SE = 0.24, p = 0.0006; contrast CS− vs. CSn, t(106) = 0.28, Estimate = 0.07, SE = 0.25, p = 0.95. On the other hand, in group 2-R no difference was found between CS+ and CS-, t(109) = −0.04, Estimate = −0.009, SE = 0.21, p = 0.99; contrast CS+ vs. CSn, t(106) = 3.09, Estimate = 0.66, SE = 0.21, p = 0.007; contrast CS− vs. CSn, t(111) = 3.01, Estimate = 0.67, SE = 0.22, p = 0.008. Figure 7

#### Unconditioned stimulus expectancy

At the end of the acquisition, participants exhibited a higher percentage of affirmative responses (YES) associated with the presentation of the CS+ for all groups, reflecting the learned contingency between the CS+ and the unconditioned stimulus during threat conditioning. The CS− and CSn stimuli, compared to the CS+, had a significantly lower probability of eliciting a YES response during acquisition (Trials 1–8).

A generalized logistic mixed-effects model including Trial as a repeated measure revealed that, during acquisition, the CS+ had higher odds of affirmative responses (YES) compared to the CS− (Odds Ratio CS+ vs CS− = 0.87, 95% CI [−0.76-98], p < 0.001) and the CSn (Odds Ratio CS+ vs CSn = 0.98, 95% CI [0.86-1.10], p < 0.001). Additionally, the unconditioned stimulus expectancy related to the CS+ remained from the end of threat acquisition (Day 1) to the assessment session (Day 3) (Odds Ratio CS+ vs CS− = 0.92, 95% CI [0.87-0.98], p < 0.001; Odds Ratio CS+ vs CSn = 0.95, 95% CI [0.90-1.01], p < 0.001). During the memory retention assessment session (Day 3), the odds of affirmative responses (YES) associated with the presentation of the CS+ remained stable (Odds Ratio CS+ vs CS− = 0.83, 95% CI [0.71-0.95], p < 0.001; Odds Ratio CS+ vs CSn = 0.90, 95% CI [0.79-1.01], p < 0.001). Supplementary material figure 3

#### Cognitive biases

##### Stimulus representation

Before conditioning (Day 1), no differences were found between groups in the levels of aversiveness for the CS+ (Planned comparison, nD2 vs. 1-R: t (137) = 0.34, p = 0.94, Estimate=0.21, SE=0.61; nD2 vs. 2-R: t (137) = 1.47, p = 0.30, Estimate=-0.84, SE=0.57; 1-R vs. 2-R: t (137) = −1.93, p = 0.13, Estimate=-1.05, SE=0.54) as well as the CS− (Planned comparison, nD2 vs. 1-R: t (137) = 1.37, p = 0.35, Estimate=0.84, SE=0.61; nD2 vs. 2-R: t (137) = −0.20, p = 0.98, Estimate=-0.12, SE=0.57; 1-R vs 2-R: t (137) = −1.76, p = 0.18, Estimate=-0.96, SE=0.54).

Differences in stimulus representation between Day 1, before training, and Day 3, after the retention test, were observed across all groups (Main Effects: Stimulus X^2^ (1,174) = 4.45, p = 0.03; Group χ^2^ (2,58) = 1.68, p = 0.19; Time X^2^ (1,174) = 1.87, p = 0.17; Conditional R^2^ = 0.523, Marginal R^2^ = 0.082).

The CS+ was scored as more aversive or unpleasant on Day 3 than on Day 1 for all experimental groups (Planned Comparison Stimulus CS+, Day 1 vs Day 3, 0-RC group: t (174) = −2.04, p = 0.04, Estimate=-0.94, SE=0.45; 1-R group: t (174) = −1.99, p = 0.04, Estimate=-0.8, SE=0.42; 2-R group: t (174) = −1.9, p = 0.05, Estimate=-0.69, SE=0.36). However, only in the 2-R group a higher rating on Day 1 was found for the CS− (Planned Comparison Stimulus CS-, Day 1 vs Day 3, 2-R group: t (174) = −1.92, p = 0.05, Estimate=0.69, SE=0.36). Figure 8a.

**Figure 8.**
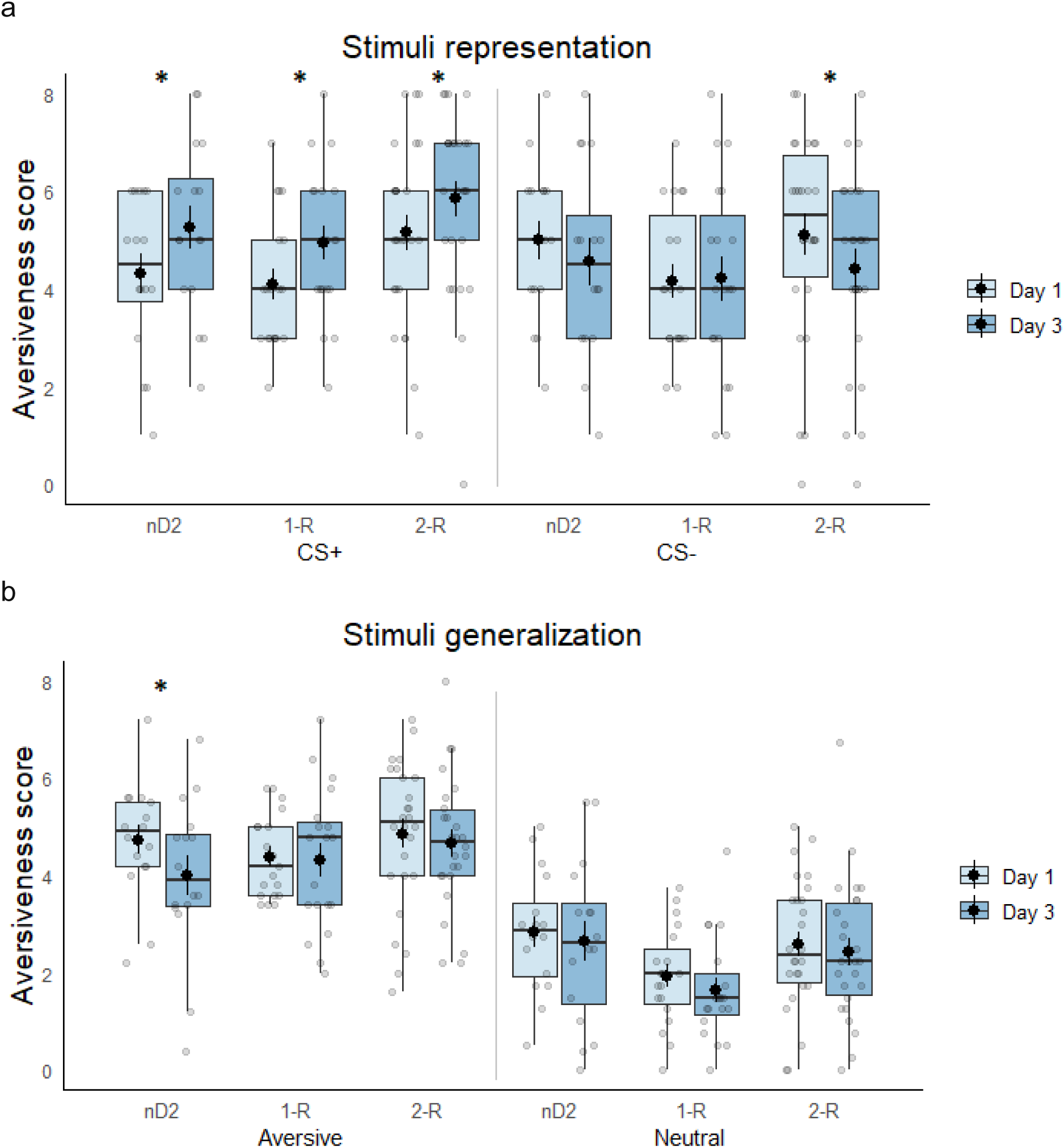
Cognitive biases assessment before threat conditioning (Day 1) and after test (Day 3). (A) Stimulus Representation (aversiveness). Aversiveness score for the CS+ and CS− on Day 1 and Day 3 for the nD2, 1-R, and 2-R groups. The boxplots indicate interquartile range (IQR), the band inside the box represents the median, and the whiskers are the maximum and minimum values. Grey circles represent the value of each individual, and a larger dark circle represents the mean. (B) Stimulus Generalization. The aversiveness score for the Aversive and neutral pictures on Day 1 and Day 3 for the nD2, 1-R, and 2-R groups. The boxplots indicate interquartile range (IQR), the band inside the box represents the median, and the whiskers are the maximum and minimum values. Grey circles represent the value of each individual, and a larger dark circle represents the mean. Asterisks (*) represent significant differences between encounters, with α=0.05

Furthermore, we evaluated all aversive and neutral faces on Days 1 and 3 for the three groups, excluding stimuli used in threat conditioning such as CS+, CS-, and CSn. For aversive faces, differences were found only in the nD2 group, showing a more aversive score on Day 1 than on Day 3 (Main Effects: Valence X^2^ (1,1028) = 417, p < 0.0001; Group χ^2^ (2,58) = 1.8, p = 0.17; Time X^2^ (1,1028) = 6.68, p = 0.009; Conditional R^2^ = 0.430, Marginal R^2^ = 0.256; Planned Comparison Aversive Stimuli, Day 1 vs Day 3, nD2 group: t (1028) = 2.68, p = 0.007, Estimate=0.72, SE=0.27). No differences were found for neutral faces for any of the groups (Planned Comparison Neutral Stimuli, Day 1 vs Day 3, nD2 group: t (1029) = −0.69, p = 0.49, Estimate=0.21, SE=0.30; 1-R group: t (1028) = 1.05, p = 0.29, Estimate=0.29, SE=0.28; 2-R group: t (1028) = 0.61, p = 0.54, Estimate=0.14, SE=0.23). Figure 8b

In Day 1 and Day 3, aversive faces were scored significantly more aversive or unpleasant than the neutral faces in all groups (Planned comparison Aversive vs. Neutral Stimuli, Day 1, nD2 group: t (1028) =6.57, p <0.0001, Estimate=1.88, SE=0.28; 1-R group: t (1028) =9.41, p < 0.0001, Estimate=2.46, SE=0.26; 2-R group: t (1028) =10.17, p < 0.0001, Estimate=2.27, SE=0.22; Day 3, nD2 group: t (1028) =4.85, p < 0.0001, Estimate=1.37, SE=0.28; 1-R group: t (1028) =10.24, p < 0.0001, Estimate=1.67, SE=0.26; 2-R group: t (1028) =9.95, p < 0.0001, Estimate=2.22, SE=0.22). Supplementary material figure 4

##### Valuation (probability and cost)

The probability of negative scenarios was higher for the CS+ than the CS− in the 1-R and 2-R groups, but no differences were found for the nD2 group. No differences were found in the probability of positive scenarios for any of the groups (Main Effects: Stimulus χ^2^ (1,2518) = 1.64, p = 0.20; Group χ^2^ (2,51) = 0.03, p = 0.97; Valence X^2^ (1,2518) = 262.5, p < 0.0001; Stimulus x Valence Interaction X^2^ (1,2518) = 5.47, p = 0.02; Conditional R^2^ = 0.220, Marginal R^2^ = 0.094; Planned Comparisons, Probability with Negative Valence, 1-R group CS+ vs. CS− t (2518) = 3.12, p = 0.001, Estimate=0.7, SE=0.2; 2-R CS+ vs. CS− t (2518) = 3.33, p = 0.0009, Estimate=0.57, SE=0.17). Figure 9a

**Figure 9.**
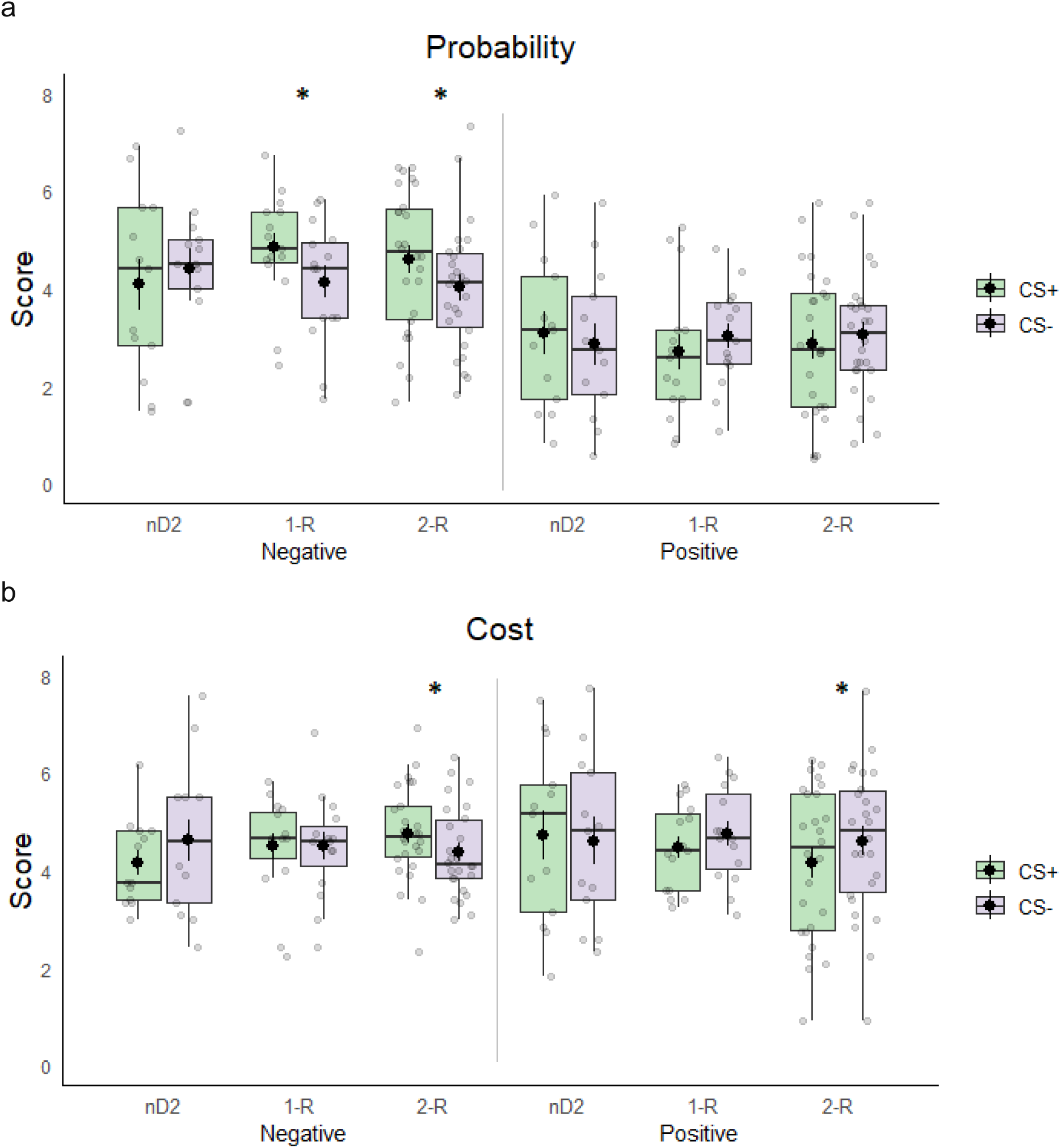
Valuation (Probability and Cost) after the test (Day 3). (A) Probability: Score for Negative and positive scenarios for CS+ and CS− in the nD2, 1-R, and 2-R groups. The boxplots indicate interquartile range (IQR), the band inside the box represents the median, and the whiskers are the maximum and minimum values. Grey circles represent the value of each individual, and larger dark circles represent the mean. A (B) Cost: Score for Negative and positive scenarios for CS+ and CS− in the nD2, 1-R and 2-R groups. The boxplots indicate interquartile range (IQR), the band inside the box represents the median, and the whiskers are the maximum and minimum values. Grey circles represent the value of each individual, and larger dark circles represent the mean. Asterisks (*) represent significant differences between encounters, with α=0.05.

When participants rated how good or bad the hypothetical situation was (positive and negative cost), no differences were found between CS+ and CS− for the 1-R and nD2 groups. However, for the 2-R group, differences were found in both positive and negative cost (Main Effects: Stimulus X^2^ (1,2518) = 1.52, p = 0.21; Group X^2^ (2,51) = 0.04, p = 0.96; Valence X^2^ (1,2518) = 0.56, p = 0.45; Conditional R^2^ = 0.229, Marginal R^2^ = 0.006; Planned Comparisons, Cost with Negative Valence, 2-R CS+ vs CS− t (2518) = 2.1, p = 0.03, Estimate=0.38, SE=0.18; Cost with Positive Valence, 2-R CS+ vs CS− t (2518) = −2.4, p = 0.01, Estimate=-0.44, SE=0.18). Figure 9b

## Discussion

Threat conditioning represents a self-preserving form of learning that fosters the creation of implicit memories and may play a role in the persistence of anxiety disorders, which is characterized by heightened reactivity to threat cues (Grupe & Nitschke, 2013). This phenomenon has been implicated in the onset of anxiety disorders (Bouton, 1993; Mineka & Kihlstrom, 1978). The impact of implicit memory on cognitive biases has been proposed as a model for various features of these disorders, such as fear incubation, difficulty of inhibited threat responses to safety cues, and stimulus generalization. Nevertheless, many experimental protocols conducted in laboratory settings often overlook the role of implicit memory in shaping cognitive biases that are essential for understanding both adaptive and maladaptive behaviors(American Psychiatric Association & American Psychiatric Association, 2013; Fernández et al., 2017; Grupe & Nitschke, 2013). Exploring the intricate relationship between implicit memories and cognitive biases may provide valuable insights into these processes, especially when considering the potentially misleading effects of examining implicit memory in isolation. Our experimental design involves manipulating implicit memory to analyze its effects on the cognitive biases associated with the stimuli used within our experimental framework to bridge this gap. Experiment 1 showed the occurrence of reminder-dependent amnesia resulting from a high-demand working memory task (HWM) performed after the presentation of the incomplete reminder. We observed a decline in implicit memory during a testing session that involved a single presentation of one conditioned stimulus (CS) from each type.

In contrast, declarative memory remained unaffected (Figures 4 and supplementary 1, respectively). Importantly, we verified that this amnesic effect on threat memory weakens biased processing toward threat stimuli. Specifically, our findings revealed that the HWM task affected implicit memory expression and altered the representation of stimuli, as evidenced by similar ratings for CS+ between Day 1 and Day 3. Conversely, the group that solely completed the HWM task on Day 2 without reminder presentation—thus lacking reactivation—exhibited higher CS+ scores on Day 3 compared to Day 1. Intriguingly, the generalization of aversiveness for angry faces showed a higher rating for this group. (Figure 5). On the contrary, the second task aimed to assess the cost and valuation of positive and negative scenarios linked to the CS+ or CS− revealed no cost-scoring disparities between negative and positive scenarios for both CS+ and CS-. Notably, higher values were observed for the CS+ in negative scenarios across both groups (Figure 6).

The current results closely align with those found in a previous report. The patterns observed in implicit and declarative memory mirror those seen during a testing session involving an extinction protocol followed by subsequent reinstatement. Thus, the incomplete reminder (the CS+ without reinforcement) effectively reactivates implicit memory, allowing the high-demand working memory (HWM) task to interfere with the implicit memory restabilization. Notably, the effects observed on stimulus representation and valuation remain consistent, irrespective of repeated presentations of various conditioned stimuli (CSs) or the unconditioned stimulus (US).

These new findings highlight the consistency in stimulus representation across experiments (Fernández et al., 2018; Picco et al., 2022), indicating a notable stability in demonstrating the aversiveness associated with implicit memory, particularly in the perception of CS+ as more aversive post-conditioning. This stability in stimulus representation validates our experimental procedures’ reliability, offering a new insight into the potential interplay between implicit memory and cognitive biases. Conversely, the assessment of probability cost and valuation yields less consistent results. This variability may stem from challenges in sustaining attention during tasks requiring the interpretation of different scenarios. Future studies aimed at simplifying the demands of these tasks could enhance their efficacy and yield more reliable outcomes with the same level of training. Another option should be observing subjective changes in these tasks when subjected to strong threat conditioning varying the number of CS+US presentations.

An additional outcome not commonly reported in post-reactivation manipulation involves the reinforcing effect of presenting a reminder without any further intervention (Forcato et al., 2014). Research on enhancing human episodic memories remains limited (Kroes et al., 2016; Forcato et al., 2016; Bavassi et al.,2019). More notably, despite the extensive use of TC in human studies to examine the reconsolidation process, there has been a significant lack of analysis concerning the effects of reactivation and reconsolidation without additional manipulation. Thus, considering our background in declarative memories (Bavassi et al., 2019; Fernández et al., 2022; Forcato et al., 2014, 2016), we designed Experiment 2 to address a specific limitation. The aim was to assess how an incomplete reminder (CS+ alone) affects implicit memory and cognitive biases. The experiment involved three groups subjected to different treatments on Day 2: no reminder (nD2), one reminder (1-R), and two reminders (2-R). The results indicated implicit memory retention for the nD2 and 1-R groups regarding the CS+ compared to the CS− and CSn. Notably, the 2-R group displayed similar skin conductance response (SCR) levels for both the CS+ and CS-, indicating a generalization effect. This effect stems more from an increase in the response to the CS− rather than a decrease in the response to the CS+ (Figure 7). This result suggests that the acquired response could generalize to a related stimulus with certain shared features, specifically the valence associated with the angry face.

In contrast, the neutral face consistently exhibited a low skin conductance response across both sessions, indicating that the generalization effect did not affect it. Consistent with prior research, the declarative component (expectancy) remained unchanged (Supplementary figure 3).

This experiment thoroughly explored the impact of incomplete reminders on cognitive biases. Our findings reveal significant differences in stimulus representation between Day 1, before training, and Day 3 across all three groups. By Day 3, the CS+ was consistently scored as more aversive or unpleasant than on Day 1 in every condition, underscoring the sustained accuracy in recognizing the aversive stimulus due to the aversive memory. Interestingly, the 2-R group scored the CS− as more aversive on Day 1 than on Day 3. We provide a plausible explanation for the observed results. This may suggest that repeated presentations of the CS+ during the reactivation session have led to enhanced accuracy in its identification. As a result, increased confidence in recognizing the reinforced stimulus indicates a growing certainty about perceiving the CS− as less aversive by Day 3 (Figure 8a). Future experiments incorporating specific confidence measures related to the responses or tasks designed to assess the accuracy of recognizing the stimuli will further support this working hypothesis.

Concerning stimulus generalization, the most notable difference observed was in the nD2 Group regarding aversive faces. This group exhibited a higher aversive score on Day 1 than on Day 3 (Figure 8b). Notably, this group did not receive any treatment on Day 2. Participants visited the laboratory for training on Day 1 and returned for testing on Day 3. We interpret this finding in light of Huppbach’s research (Hupbach et al., 2008), which shows that the spatial context plays a unique role in episodic memory updating. The same spatial context is essential for memory malleability during both original and new learning, alongside presenting a specific reminder of the target memory. Based on this premise, returning to the laboratory may act as a partial reminder, which was missing in the nD2 Group, potentially explaining the differences in results across the various measures of this protocol.

The findings from the probability and cost task designed to assess cognitive biases suggest that the probability of negative scenarios was higher for the CS+ than the CS− in the 1-R and 2-R groups; however, no significant differences were observed in the nD2 Group. This absence of variation within the nD2 Group further supports the notion of diminished accuracy related to the protocol that did not involve any manipulations on Day 2. Additionally, consistent with the increase in accuracy associated with the potential degree of reactivation, noticeable differences emerged in the 2-R Group when participants rated negative costs. In contrast, the 1-R and nD2 groups showed no such effects. This outcome may be understood as a consequence of the repeated reactivation manipulation, which served to enhance the attributes of the reactivated memory multiple times (Forcato et al., 2014).

Furthermore, in alignment with the results observed for stimulus representation, the 2-R group rated scenarios linked to the CS− as more positive than those associated with the CS+, thereby reinforcing the interpretation of stronger confidence in the less aversive nature of the CS-. We might also consider that within this paradigm, the spatial context—specifically the laboratory—contributes significantly, as suggested for the nD2 Group. However, merely having this context is insufficient to account for memory malleability, as it requires an incomplete reminder, the CS+ in the absence of reinforcement, as the experimental manipulation serves as a crucial element for memory reactivation.

The findings from both experiments examining the generalization analysis for the stimulus representation task should indicate increasing maintenance of aversiveness when participants are exposed to incomplete reminders. The absence of treatment on Day 2 suggests a loss of aversiveness specificity to the CS+ in the nD2 Group, while arriving at the laboratory without prior reminder exposure has the opposite effect. Overall, groups exposed to one or two reminders (R) can maintain the aversiveness associated with the CS+ and do not generalize that aversiveness to other angry faces.

The findings related to the 2-R Group warrant a more thorough examination. The experimental design, which explored implicit memory and cognitive biases, enabled us to identify a specific time at which implicit memory changes do not influence the valuation of stimuli. Our findings indicated a generalization effect regarding implicit memory, as participants exhibited comparable skin conductance response (SCR) levels for the CS+ and CS− on Day 3. Additionally, there was a noticeable specificity of stimuli related to cognitive bias, evidenced by increased levels of aversiveness for the conditioned stimulus CS+, and a reduction in aversiveness for the conditioned stimulus CS-. This disparity may be related to the dissociation observed in certain aspects of fear memory expression, particularly as described by Kindt’s laboratory (Kindt et al., 2009; Soeter & Kindt, 2010). The researchers highlighted the different dynamics involved in updating implicit and declarative memories, noting that the latter is influenced by cognitive biases stemming from the former. The initial report (Soeter & Kindt, 2015) focused on memories associated with spider phobias. In this study, participants who feared spiders received 40 mg of the noradrenergic β-blocker propranolol following a brief exposure to a tarantula. This intervention disrupted the reconsolidation of fear memory, which led to a transformation of avoidance behavior into approach behavior. However, there was no immediate effect on the participants’ self-reported fear of spiders. Remarkably, the alignment between behavior and declarative memory became apparent three months later. Building upon these findings, the researchers conducted another experiment to analyze the dynamics of both memory types with the same intervention. In this case, the first behavioral test was conducted two days or four weeks after the administration of propranolol. They described a reduction in spider avoidance behavior and self-reported fear over the course of a year, particularly with the earlier testing. Importantly, they replicated the time-dependent difference in the onset of the declarative components (Peters et al., 2025). This dissociation is also evident in another report (Andreatta et al., 2015), where participants were guided through two virtual office contexts (CSs) in which electric shocks (US) were unpredictably administered in one of the settings. The testing session incorporated a generalization context that featured elements from reinforced and unreinforced CSs. The results indicated a heightened startle response to the CS+ compared to the CS-, with the CS+ perceived as more aversive. Surprisingly, implicit and explicit responses were dissociated within the generalization context; it was rated as more negative despite showing a startle response similar to the CS-. The authors suggest that this dissociation may indicate distinct and specific generalization processes. Considering this background, future experiments may include a second test, considering an intersession interval of around 30 days.

Finally, we underscore the experimental value of this protocol by combining the TC memory with its impact on cognitive biases. First, actual outcomes are consistent with our prior research, supporting that reminders featuring harmful prediction errors, marked by the absence of congruence between the training trial and reactivation session, due to the lack of the US, destabilize consolidated memory given the opportunity to modify it (Fernández et al., 2016; Sevenster et al., 2013). These modifications might mirror the effect on cognitive biases (Experiment 1) in some cases but not others (Experiment 2), revealing different processing paths for implicit memory and the valuation of the stimuli associated with it. Second, in this report, we analyze a 3-day memory. A question emerges considering when we consider older memories. In this sense, it would be possible that the passage of time determines a system consolidation. At this point, the TC memory manipulation did not affect the acquired cognitive biases. Moreover, it would also be possible that the simple passage of time determines the loss of TC memory expression despite the maintained cognitive biases, showing how low or absent TC memory could coexist with cognitive biases. Ultimately, this experimental approach should be considered as another tool for deepening our understanding of the impact of post-reactivation interventions, potentially opening avenues for novel treatments for maladaptive memories (Elsey et al., 2018; Kroes et al., 2016; Lane et al., 2014; Schroyens et al., 2023).

