## Supplementary material for "Reactivation of threat conditioning memory in humans: disentangling the effects on emotional memory and cognitive biases"

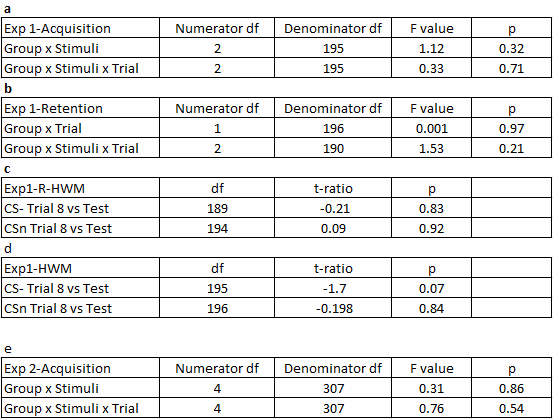

Supplementary table 1. a,b,c) Complementary results of the Experiment 1 and e) Complementary results of the Experiment 2.

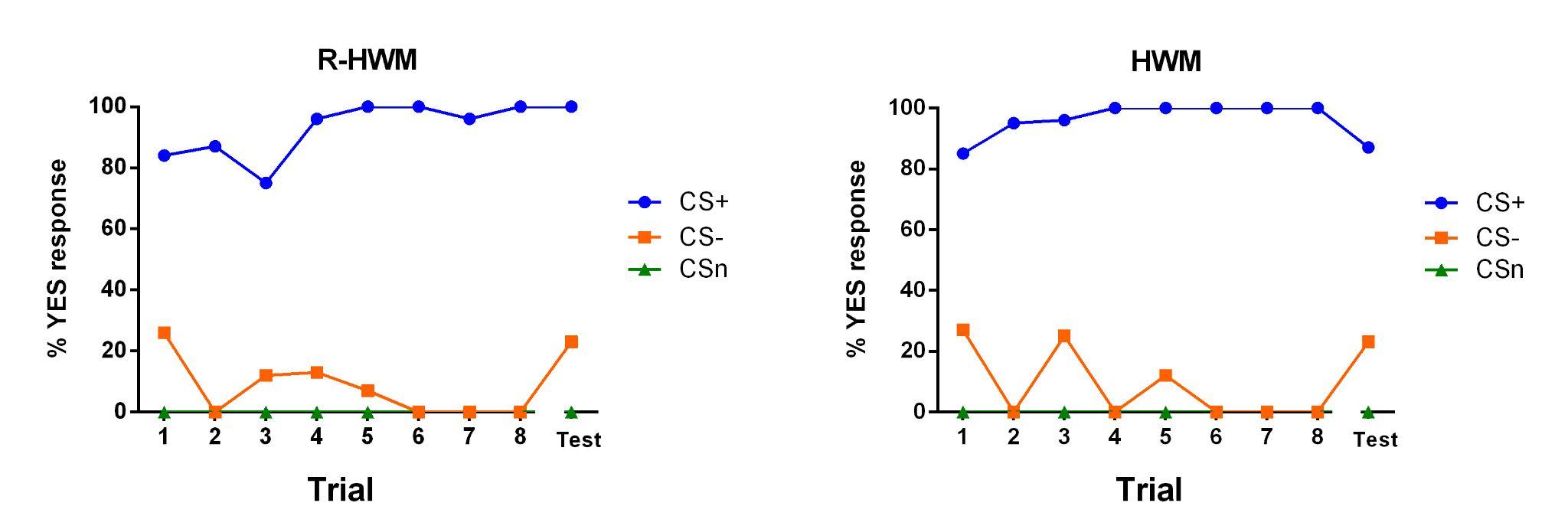

Supplementary figure 1. Percentage of YES responses for each stimulus (CS+, CS-, and CSN) for the 8 trials on Day 1, and the trial on Day 3 (testing), for the R-HWM and HWM groups.

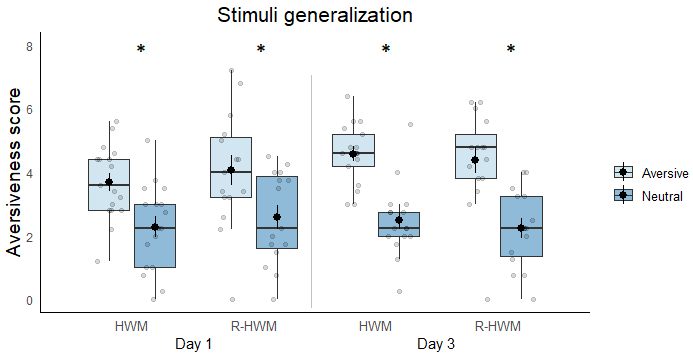

Supplementary figure 2. Stimulus generalization. Aversiveness score for all aversive and neutral faces (excluding those used in conditioning), for Day 1 and Day 3 in both groups (HWM and R-HWM).

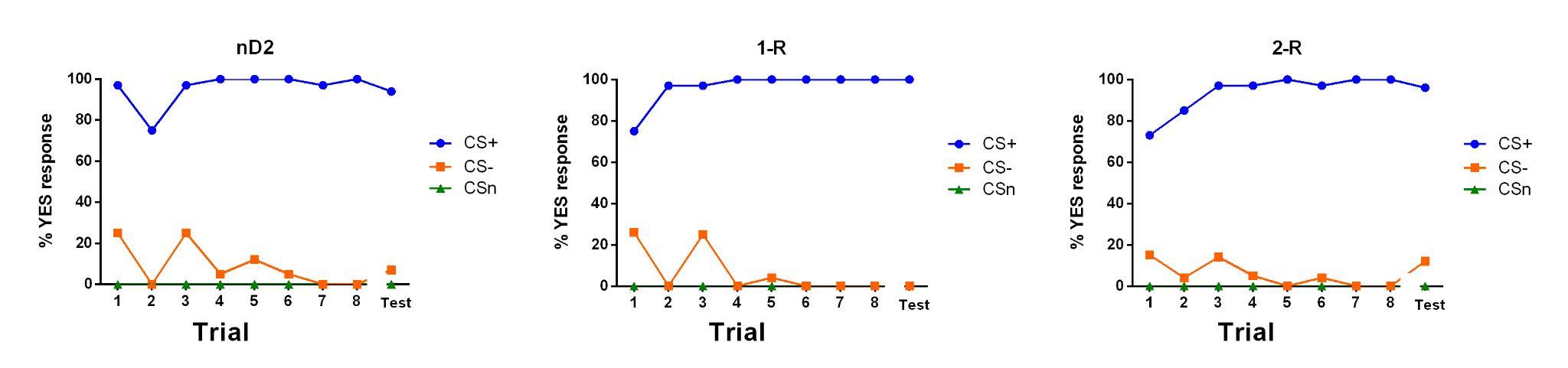

Supplementary figure 3. Percentage of YES responses for each stimulus (CS+, CS-, and CSN) for the 8 trials on Day 1, and the trial on Day 3 (testing), for the nD2, 1-R and 2-R groups.

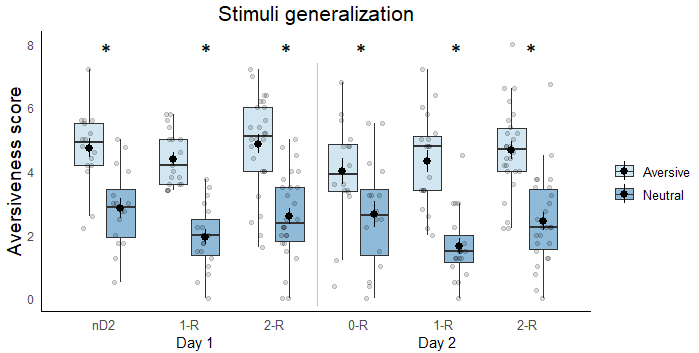

Supplementary figure 4. Stimulus generalization. Aversiveness score for all aversive and neutral faces (excluding those used in conditioning), for Day 1 and Day 3 or the nD2, 1-R and 2-R groups.
